# *REVEILLE2* Thermosensitive Splicing: A Molecular Basis for the Integration of Nocturnal Temperature Information by the Arabidopsis Circadian Clock

**DOI:** 10.1101/2023.04.24.538045

**Authors:** Allan B. James, Chantal Sharples, Janet Laird, Emily May Armstrong, Wenbin Guo, Nikoleta Tzioutziou, Runxuan Zhang, John W.S. Brown, Hugh G. Nimmo, Matthew A. Jones

## Abstract

Cold stress is one of the major environmental factors that limit growth and yield of plants. However, it is still not fully understood how plants account for daily temperature fluctuations, nor how these temperature changes are integrated with other regulatory systems such as the circadian clock.
We demonstrate that REVEILLE2, a MYB-like transcription factor, exhibits a cold-induced alternative splicing switch from a non-translatable isoform at ambient temperature to a translatable isoform upon cold exposure. We explore the biological function of *REVEILLE2* using a combination of molecular genetics, transcriptomics, and physiology.
Disruption of the *REVEILLE2* cooling switch alters regulatory gene expression, impairs circadian timing, and improves photosynthetic capacity. Changes in nuclear gene expression are particularly apparent in the initial hours following chilling, with chloroplast gene expression subsequently up-regulated.
The *REVEILLE2* cold switch extends our understanding of plants immediate response to cooling. We propose that the circadian component *REVEILLE2* restricts plants responses to nocturnal reductions in temperature, thereby enabling appropriate responses to daily environmental changes.

**Plain language summary:** Plants need to respond appropriately to temperature, accounting for the expected daily patterns of reduced temperatures that occur every night relative to the day. Here, we show that a gene expressed at night fulfils this function.

## Introduction

Circadian clocks have evolved in most organisms as a response to the daily rotation of the Earth and the imposition of alterations in light:dark and temperature. Plants gain a fitness advantage by aligning internal processes to their external environment (Dodd *et al*., 2005) – the clock oscillator mediates this harmonious relationship (Nagel & Kay, 2012; Hsu & Harmer, 2014). While temperature changes associated with daily light:dark cycles are inevitable, temperature in nature fluctuates rapidly and dynamically across timescales. Temperature is also a key seasonal indicator, such that many plant species need to experience the chill of winter to permit flowering in the spring (Penfield, 2008; Andres & Coupland, 2012).

Although specific temperature sensors are being described (Jung *et al*., 2016; Chung *et al*., 2020; Jung *et al*., 2020; Antoniou-Kourounioti *et al*., 2021), it is also apparent that temperature sensing is broadly distributed across multiple regulatory networks, including those routed through the circadian clock (Kidokoro *et al*., 2017; Antoniou-Kourounioti *et al*., 2018; Calixto *et al*., 2018). The relationship between the clock and temperature is complex, but important on a number of levels including; 1) the clock is temperature compensated, whereby the pace of clock oscillations is relatively stable across a wide range of physiologically relevant temperatures (Hastings & Sweeney, 1957; Pittendrigh 1954); 2) the clock can be entrained by small differences in temperature cycles (Zimmerman *et al*. 1968; Rensing & Ruoff, 2002), but 3) the clock becomes somewhat damped following several days of low temperature (Ramos *et al*., 2005; Bieniawska *et al*., 2008). Within minutes of exposing Arabidopsis plants to low temperature, changes in transcript accumulation can be detected followed by waves of changes in transcriptome composition (Fowler & Thomashow, 2002; Maruyama *et al*., 2004; Vogel *et al*., 2005). By 24h the transcript levels of more than 1000 genes have either increased or decreased, with the induction of the *C-REPEAT-BINDING FACTOR (CBF)1*,*-2* and *-3* transcriptional activator genes (also known as *DEHYDRATION-RESPONSIVE ELEMENT-BINDING FACTOR (DREB)1b*, *-1c* and *-1a*, respectively) occurring within 15 min of low temperature exposure (Chinnusamy *et al*., 2007). Cold induction of the *CBFs* is gated by the circadian clock; their expression is dependent on the time of day that the plants are exposed to low temperature with expression more pronounced around subjective dawn compared to subjective dusk under constant light conditions (Fowler *et al*., 2005).

In *Arabidopsis thaliana* the core circadian oscillator features multiple interlocking feedback loops of gene expression and comprises dawn-expressed CIRCADIAN CLOCK ASSOCIATED 1 (CCA1) and LATE ELONGATED HYPOCOTYL (LHY) and the evening phased transcriptional repressor PSEUDO-RESPONSE REGULATOR 1 (PRR1, also known as TIMING OF CAB EXPRESSION 1, TOC1). Additional components of the clock are arranged in a complex set of gene expression loops with multiple interactions; the principal components include day-phased transcriptional repressors PRR9, PRR7 and PRR5, and the evening-phased components EARLY FLOWERING 3 (ELF3), ELF4 and LUX ARRHYTHMO (LUX) which interact to form a transcriptional repressor named the Evening Complex (EC).

LHY and CCA1 belong to the REVEILLE (RVE) family of proteins, all of which contain a single DNA-binding Myb-like domain. Other family members include RVE1 through RVE8, in addition to a RVE7-like protein. RVEs appear to play contrasting roles in controlling the pace of the circadian clock; mutations in *CCA1* and *LHY* shorten clock pace (Schaffer *et al*., 1998; Wang & Tobin, 1998; Green & Tobin, 1999; Mizoguchi *et al*., 2002), while mutations in *RVE8*, *6* and *4* lengthen the circadian period (Farinas & Mas, 2011; Rawat *et al*., 2011; Hsu *et al*., 2013). RVE5 and RVE3 display more subtle effects on setting the tempo of the clock (Gray *et al*., 2017), and RVE1 does not affect clock pace but instead mediates the circadian regulation of the auxin pathway (Rawat *et al*., 2009). Constitutive expression of either RVE2 or RVE7 has previously been reported to repress circadian gene expression (Kuno *et al*., 2003; Zhang *et al*., 2007). More recently, work has revealed that RVE4 and RVE8 contribute to cold-induced gene expression, demonstrating how RVE proteins contribute towards the integration of temperature signals into the circadian system (Kidokoro *et al*. 2021, 2023).

Clearly there is a close relationship between temperature and plants’ timing mechanism. However, how very early temperature perception is transduced to the clock, and how the circadian system regulates (or ‘gates’) clock output pathways is not fully understood (Panter *et al*., 2019). Towards addressing these questions, work employing temperature transitions (James *et al*., 2012a; James *et al*., 2012b; Calixto *et al*., 2018) and natural temperature fluctuating environments (Annunziata *et al*., 2018; Matsubara, 2018) has greatly extended our understanding of plants’ responses to temperature and demonstrate the role of rapid and dynamic reprogramming of the transcriptome (Calixto *et al*., 2018), driven by alternative splicing (AS), as central to plants’ plastic responses to cooling. A striking feature of the changes occurring at the AS level with cooling is the induction of transcript isoform switches (Guo *et al*., 2017; Calixto *et al*., 2018; James *et al*., 2018). Here the ratios of specific transcript isoforms flip rapidly and reversibly with temperature change. For example, isoform switching of transcript isoforms of *LHY* involves either the canonical splicing or retention of the first intron in the 5’UTR. This switch is sensitive to temperature reductions as small as 2°C and is scalable over a wide dynamic range of temperature (James *et al*., 2018). It is therefore increasingly apparent that alternative splicing of transcripts can be utilised to rapidly respond to environmental changes (Laloum *et al*. 2018).

Our previous work employing ultra-deep RNA-sequencing (RNA-seq) of a diel time-course of plants exposed to low temperature showed that around 900 isoform switches are engaged upon cooling, and around 500 of these occur within 6h (Calixto *et al*., 2018). Here, we extend this work to characterise the biological role of an isoform switch within the *RVE2* transcript that is induced within 20 minutes of chilling. A T-DNA insertional mutant (*rve2-2*) that is devoid of cold-switched *RVE2* displays increased accumulation of key regulatory transcripts including *CBFs*. *rve2-2* seedlings have impaired circadian rhythms at reduced temperatures and temporal profiling shows most Differentially-Expressed Genes (DEGs) comprise a wave of gene expression initiated shortly after peak RVE2 expression during transient cool nights. However, we also identified a group of chloroplast-encoded genes that are repressed in *rve2* plants following the initial DEGs which is correlated with altered photosynthetic capacity the following morning. Our data therefore suggest that *RVE2* represses the induction of low temperature responses during the night to enable an appropriate photosynthetic and cold-response strategy with the promise of the ensuing dawn.

## Materials and Methods

### Plant material and growth conditions

All plant material was the Columbia (Col-0) ecotype. The *rve2-2* mutant was obtained from GABI-Kat (Kleinboelting *et al*., 2012) and homozygous mutant lines selected and characterised (Suppl Fig. S3). The single T-DNA insertion lines *ptb1-1* (SALK_013673C) and *ptb2-*1 (SAIL_736_B12) and the amiRNA knockdown line ami*PTB1&2* (*ami1-1;2-1*; Ruhl *et al*., 2012) were a gift of Prof. Dr. Andreas Wachter (Johannes Gutenberg University, Mainz). The single T-DNA insertion lines *mkk1-2* (SALK_027645) and *mkk2-1* (SAIL_511_H01) (Gao *et al*., 2008) were a gift of Prof. Heribert Hirt (King Abdullah University of Science and Technology). *r*ve2-2 *CCA1::LUC2* lines were generated by crossing *rve2-2* with a Col-0 reporter line (Jones *et* al. 2015). The *RVE2::RVE2:LUC* construct was generated by cloning the *RVE2* locus (including 1000bp upstream of the start codon) into pDONR221 before being transferred into pGWB535 using Gateway cloning (Invitrogen). *RVE2::RVE2:LUC* was transformed into *rve2-2* seedlings using agrobacteria-mediated transformation (Clough and Bent 1998). Growth conditions for the RNA-seq diel and temperature time series experiments are detailed in the Extended Materials and Methods section of the Supporting Information Document.

### DNA & RNA extraction, cDNA synthesis, RT-PCR, and qPCR

RNA extraction, cDNA synthesis, qPCR (RT-qPCR) and High resolution (HR) RT-PCR were performed as described in the Extended Materials and Methods section of the Supporting Information Document in line with previous work (Simpson *et al*., 2008; James *et al*., 2012a; Simpson *et al*., 2016; James *et al*., 2018). Primer sequences are provided in Suppl Table S1.

### RNA-seq

RNA-seq libraries were constructed using the Illumina TruSeq library preparation protocol. Libraries had an average insert size of 280 bp and were sequenced three times on the Illumina HiSeq 2500 platform to generate 100 bp paired end reads. After standard quality control and trimming of reads, transcript expression was determined using Salmon version 0.82 (Patro *et al*., 2017) in conjunction with AtRTD2-QUASI (Zhang *et al*., 2017).

### Chlorophyll fluorescence

Chlorophyll fluorescence parameters were recorded with an IMAGING-PAM Maxi chlorophyll fluorescence system (Walz). Approximately 30 individually spaced seedlings were entrained for 12 days in 12:12 light:dark cycles on half-strength Murashige and Skoog (MS) medium without supplemental sucrose for 12 days before transfer to the imaging chamber. After transfer from the growth chamber plants were dark adapted for 30 min prior to determination of *F_v_/F_m_* at ZT2. At the following dusk, plants were chilled to 4°C, and then maintained at 4°C for the remainder of the experiment. Plants were dark-adapted for 30 min prior to determination of *F_v_/F_m_* at ZT2.

### Luciferase imaging

Plants were entrained for 6 days in 12:12 L/D cycles under white light on MS medium without sucrose before being sprayed with 3-mM D-luciferin in 0.01% (v/v) Triton X-100 as previously described (Battle & Jones, 2020). Imaging was completed over 5 days using a Teledyne Photometrics LUMO camera controlled by μManager (Edelstein et al., 2014) before data were processed using ImageJ (Schneider et al., 2012). Patterns of luciferase activity were fitted to cosine waves using fast Fourier transform–nonlinear least squares to estimate circadian period length (Zielinski et al., 2014).

### Statistical methods

The RNA-seq data quality was checked with principal component analysis (PCA) plot and sample distribution plot by using the 3D RNA-seq App (Guo et al., 2021) (Extended Materials and Methods). Differential expression analysis of the RNA-seq data focussed on the day 2 transient-cooling data. For this dataset low expressed genes (those with a cumulative TPM <80 across the time-points for day 2) were filtered to increase statistical power. Transcript isoforms were considered differentially expressed if the difference between Col-0 and rve2-2 was at least 1.2-fold for 3 contiguous time points. Analysis of variance (ANOVA) was used to test the variance in the RNA-seq TPM values on the explanatory factors genotype and time. The Anova function in the ‘car’ (Companion to Applied Regression) package (version 3.1-1) in R was used to prepare the type-II ANOVA tables for the differentially expressed candidates (Suppl Dataset S5). Fisher’s exact test for testing the null of independence of rows and columns in a contingency table with fixed marginals were performed in R using the fisher.test function in the ‘stats’ package (The R Stats Package, version 4.2.2). Other analyses and visualisations of data were carried out using the ggplot2 package (version 3.4.1; (Wickham, 2016)) in R version 4.2.2 (2022-10-31). Hutcheson’s t test was used to compare differences in Shannon’s diversity index (Hutcheson 1970).

### GO term enrichment analysis

The Gene Ontology (GO) over-representation test (Boyle *et al*., 2004) was implemented in clusterProfiler(Yu *et al*., 2012) using the parameters pAdjustMethod = “BH”, pvalueCutoff = 0.05, qvalueCutoff = 0.1.

## Results

### Nocturnal chilling induces rapid changes in patterns of *RVE2* alternative splicing

We previously identified thousands of genes which were differentially expressed (DE) and/or differentially alternatively spliced (DAS) in a high-resolution RNA-seq time series experiment of Arabidopsis plants exposed to low temperature (Calixto *et al*., 2018). Genes can be regulated by both transcription and AS (DE + DAS); around 800 of these were identified by Calixto et al., 2018. Some genes with large Δ percent spliced (ΔPS) can have low expression levels and others with lower ΔPS can have high expression levels. To ascertain which DE + DAS genes merited further characterisation we plotted the ΔPS spliced ratio with the equivalent gene expression level for around 35k transcripts from genes expressing at least two different transcripts. We found that *RVE2* (*At5g37260*, also known as *CIRCADIAN1 [CIR1;* (Zhang *et al*., 2007)] has the largest combined change in DE and DAS (Fig. **1a** and Supplementary Dataset **S1**).

**Fig. 1.**
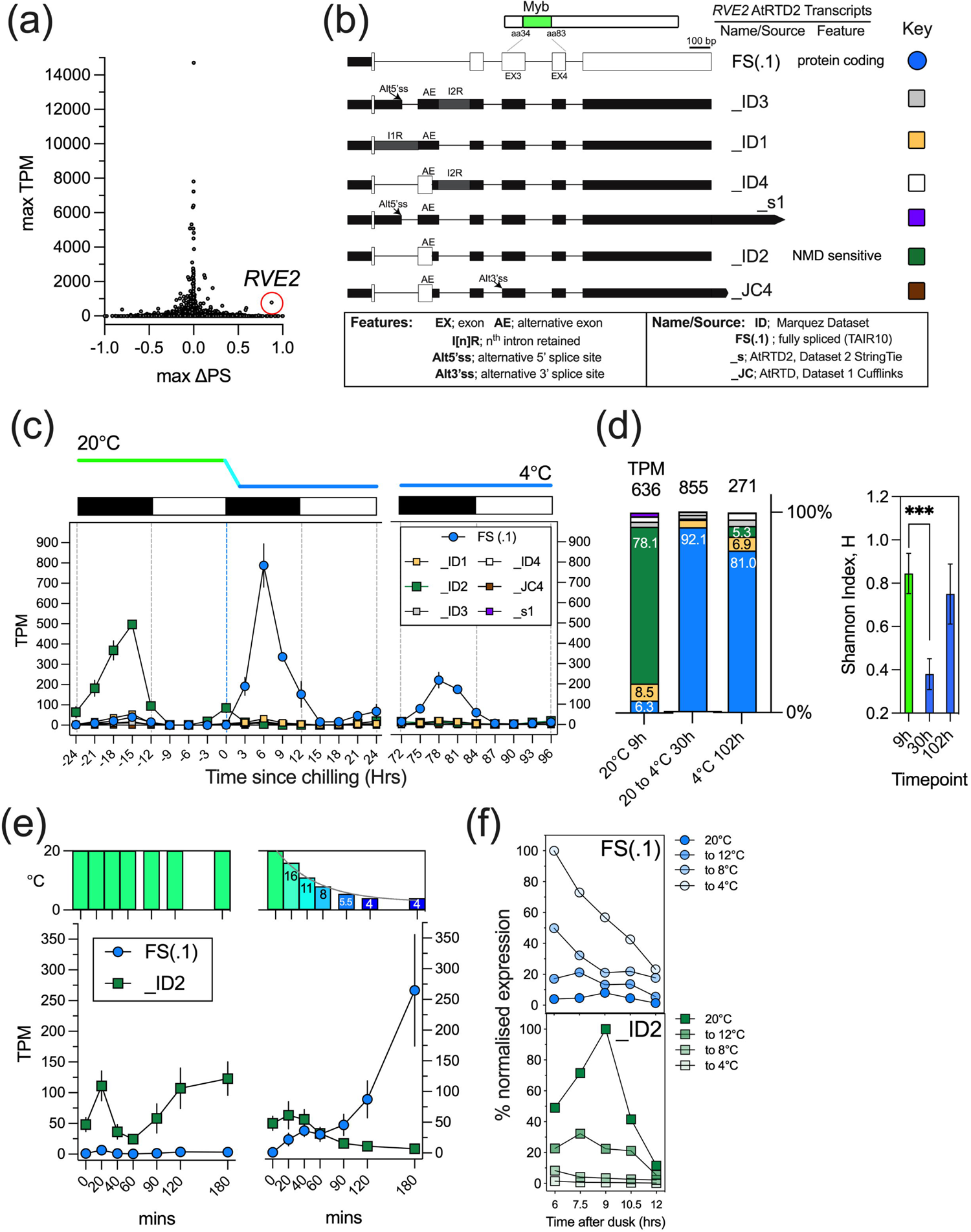
RNA-seq reveals altered patterns of alternative splicing during chilling. (a) Relationship between Δ percent spliced (ΔPS) and expression level (TPM) for 35k transcripts for genes expressing at least two transcripts, Supplementary Dataset **S1**. Red circle denotes *RVE2* FS (.1). (b) Diversity of *RVE2* transcripts. Transcript maps of seven differentially spliced *RVE2* transcripts identified using AtRTD2-QUASI (central boxed area labelled ‘RTD2’), including the FS.1 (AT5G37260.1) and AE (AT5G37260_ID2) variants. TAIR10 and Araport11 include only the FS (.1) transcript isoform. Transcript maps: white boxes – coding sequences; black boxes – UTRs; grey box - intron retention. Where an AS event causes introduction of a premature stop codon and loss of open reading frame, the downstream region is represented by black boxes (UTRs). (c) Temperature and diel time series RNA-seq profile of the cold-induced *RVE2* isoform switch (mean and ± SEM [*n*=3] of normalised expression levels [TPM; Transcripts Per Million]) for the denoted transcript isoforms. Data is plotted relative to the initiation of chilling to 4°C (dashed blue vertical line), diurnal 12h dark/12h light conditions as black/white rectangles, respectively. (d) Left Stacked bar representation of % abundance of denoted isoforms at 20°C (_ID2 peak at 9h timepoint), day 1 transition to 4°C and day 5 acclimation to 4°C (FS .1) peaks at the 30 and 102h timepoints, respectively); total TPM for timepoints denoted above the stacked bars, representative abundances (%) denoted within bars or to side of bars. Right Diversity of transcript isoforms at each time point is quantified using the Shannon Index. (e) *RVE2* cold-switching starts at onset of cooling. *RVE2* transcript isoform expression data from IE RNA-seq dataset (Supplementary Dataset **S2**) and is mean and ± SEM (*n*=4) of expression levels (TPM; Transcripts Per Million) for the FS (.1) and _ID2 isoforms for the first 3h of the dark period, either for plants experiencing no cooling (*left*, steady state 20°C), or for plants subjected to cooling (*right*, cooling transition). Temperature bar (top) indicates growth cabinet temperature at the denoted time points. (f) The *RVE2* cold-switch in plants under-going different extents of cooling was examined using RT-qPCR, *upper* FS(.1) isoform and *left* AE isoform. Plants were entrained in 20°C in LD cycles then subjected to the denoted cooling extent at the onset of dusk with samples harvested at time-points spanning mid-dusk to dawn. The ΔΔCt method was used to calibrate normalised Ct values (using the average of Cts for IPP2 and ISU1 housekeeping genes) to the time-point demonstrating maximal expression (6h at 20 to 4°C for FS(.1), upper; and 9h at 20°C for AE, *lower*).

*RVE2* has several detectable AS events including the inclusion of an alternative exon (AE) in intron 1 (Fig. **1b**). Analysis of RNA-seq data (Calixto et al., 2018) with Salmon/RTD2-QUASI (Zhang et al., 2017) identified seven transcripts containing various combinations of these AS events. The FS(.1) isoform encodes the full-length protein with a single MYB domain (Fig. **1b**), while the other variants are likely non-functional since they encode transcripts with a premature termination codon after either 4 or 23 amino acids (PTC; Fig. **1b**, **1c**, Suppl Fig. **S1a**; Raxwal et al., 2020). Assessment of *RVE2* AS during a diel time-course including a period of chilling shows that at 20°C *RVE2* transcript accumulation consists almost entirely of the non-productive isoforms (green line, Fig. **1c**) whereas for plants undergoing cooling and for plants acclimated to 4°C, the *RVE2* transcript population primarily comprises the protein-coding FS(.1) isoform (blue line, Fig. **1c**, Suppl Fig. **S1a**; Calixto et al., 2018). At the peak of *RVE2* expression at 20°C FS(.1) constitutes only 6.3% of the total gene expression, whereas _ID2 is 78.1% (9h data, Fig. **1d**, Shannon diversity index = 0.84). Cold switching increases the uniformity of *RVE2* alternative splicing outcomes and raises FS to 92.1% of total abundance at gene level while reducing _ID2 to 0.6% (30h data, Fig. **1d**, Shannon diversity index = 0.38; p < 0.001). Following an extended period of cooling, peak *RVE2* expression is damped but FS(.1) remains the major component at 81% (102h data, Fig. **1d**, Shannon diversity index = 0.75). The _ID1 isoform, comprising intron 1 as well as the AE (IR1 + AE) was evident in the 20°C and 4°C acclimated peaks, albeit at low levels (8.5 and 6.9%, respectively, Fig. **1d**) and therefore the *RVE2* cold switch primarily featured the flipping in abundance of the FS(.1) and _ID2 isoforms (Suppl Fig. **S1a**). Calixto et al., 2018 listed *RVE2* within a group of 59 TF genes that were DAS within 0-6h of cooling (Supplemental Dataset 12C of Calixto et al., 2018). We refined further the rapid induction of the *RVE2* isoform switch by capturing ‘immediate-early’ (IE) responding genes in an RNA-seq experiment for cooled Arabidopsis plants focussed on the first 3h of cooling (Fig. **1e**, Suppl Dataset **S2**). For this, we initiated cooling at dusk, consistent with the experiment of Fig. **1c**. The data show that the *RVE2* isoform switch is induced within 20 min of exposure to cooling; FS(.1) levels rise, and _ID2 levels dip relative to the equivalent time points at 20°C (Fig. **1e**). Within this timeframe the temperature of the growth chamber had dropped to 16°C (Fig. **1e**). By 90 mins of cooling _ID2 levels were negligible, and there was a clear upswing in FS(.1) from 20 min onwards (Fig. **1e**). Analysis of 1,286 genes that are DAS across the first 3h of cooling in the IE RNA-seq dataset shows that *RVE2* is one of only two genes that are DAS within the first 20 min (Suppl Fig. **S1b**). To validate this, we performed isoform specific RT-qPCR for time points immediately after the onset of cooling (Suppl Fig. **S1c**). These data confirm the rapid cold-switch; transcripts containing the Alternative Exon (AE) were lower after only 15min of cooling, whereas FS(.1) was higher after 60min of cooling (Suppl Fig. **S1d**). Student’s t-tests were performed to compare expression levels of FS(.1) and AE between the 20°C and the cooling to 4°C treatments. There was a significant difference in AE levels at 15 mins between the 20°C treatment and the cooling treatment (p = .0004). There was also a significant difference in FS(.1) levels at 60 mins between the 20°C treatment and the cooling treatment (p = < .0001). We examined the dynamics of the cold-switch for plants under-going different extents of cooling. Here we found that the *RVE2* switch is sensitive and scalable with the temperature drops that we employed (from Δ8 to Δ16°C; Fig. **1f**).

### Patterns of *RVE2* alternative splicing are maintained for several days following chilling

Up until this point we had challenged plants with dynamic cooling (transition, and then adaption, to low temperature) whilst maintaining predictable alterations in light and dark phases. We next asked whether the cooling associated emergence of the FS(.1) oscillation persisted in the absence of these diurnal light:dark (LD) cues. Initial experiments using isoform specific RT-qPCR showed that *RVE2* rhythms persisted under 12°C LL conditions (Suppl. Fig. **S2**). The FS(.1) isoform comprised around 15% of total transcript levels at 20°C in LD (Suppl. Fig. **S2a**), and while these levels reduced further in LL at 20°C, low amplitude rhythms of FS(.1) emerged during the first subjective dark phase of the 12°C LL free run (Suppl Fig. **S2b**). Oscillations of the AE isoform persisted in 20°C LL but became less rhythmic in LL at 12°C (RAE>0.6, Suppl Fig. **S2c**). The estimated period lengths appeared distinct for the AE and FS(.1) isoforms at 20°C LL (∼23h for AE in 20°C LL and ∼22h for FS(.1), Suppl Fig. **S2b**), although further work will be needed over extended timescales to obtain more reliable period estimates.

We extended these observations by using RNA-seq (Fig. 2, Suppl Dataset **S3**) to concomitantly monitor of all seven isoforms over a period of 56h in LL. Like the equivalent RT-qPCR data in Suppl Fig **S2** we found that the _ID2 isoform maintained a robust (higher amplitude) rhythm in 20°C LL (Fig. **2a**, **c**) compared to the severely damped FS(.1) oscillation in 20°C LL (Fig. **2a**, **b**). By contrast, the _ID1 and _ID4 expression levels were relatively constant although we did observe a change in phase in both isoforms following chilling (Fig. **2d**, **e**).

**Figure 2.**
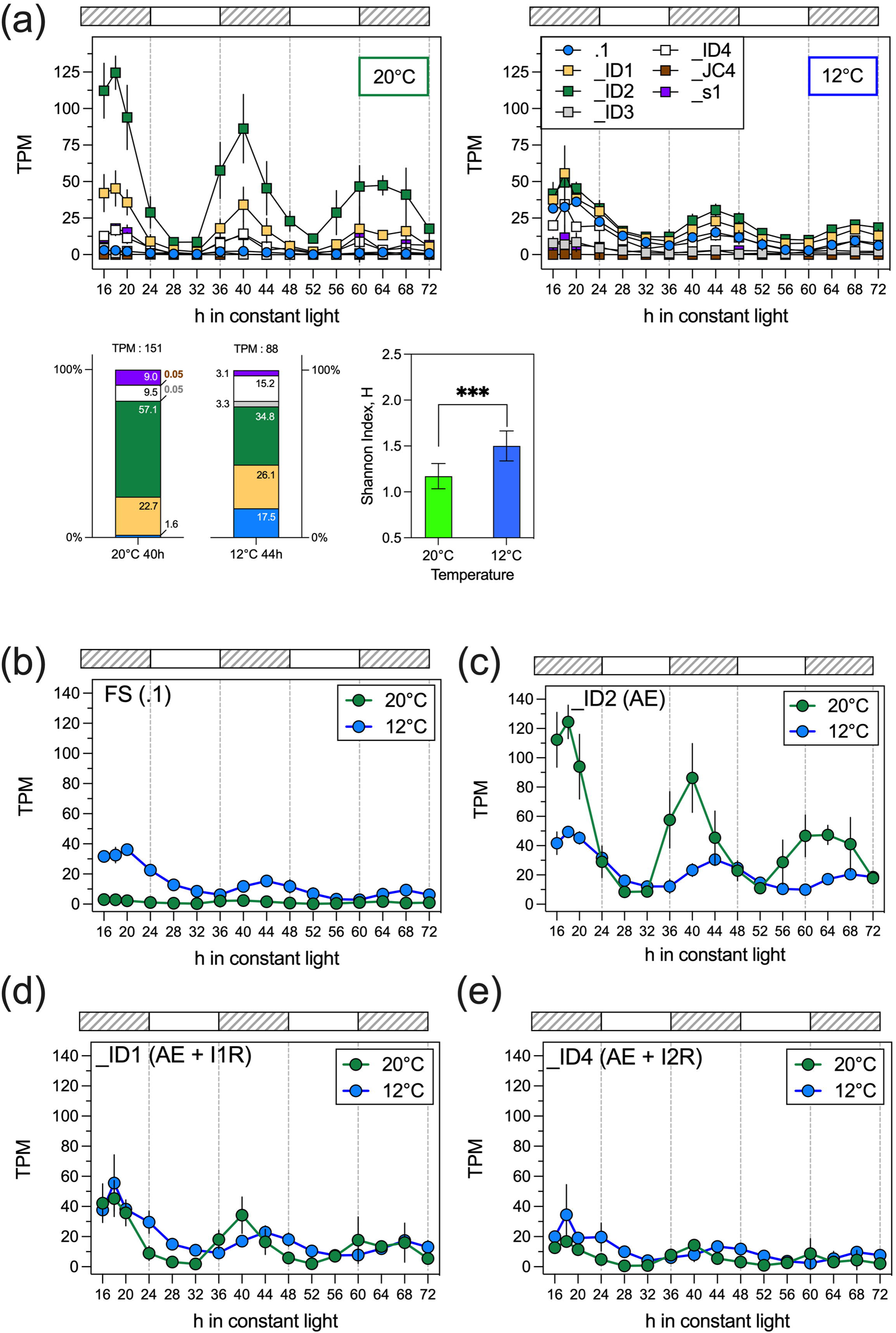
Alternative Splicing of *RVE2* persists in free-run conditions. Isoform specific expression of *RVE2* in constant light (LL) at either 20°C or 12°C by RNA-seq (Supplementary Dataset **S3**). Col-0 plants were grown to maturity (5 weeks) in 12h dark: 12h light at 20°C, then either held at 20°C or chilled to 12°C during the final night prior to release into LL for 56h. Subjective dark phases as light-grey shaded boxes. Plants were harvested every 4h, except an additional time point at 18h. (a) Profiles for the denoted transcript isoforms; 20°C LL, *upper left* graph, 12°C LL, *upper right* graph. *Lower left* graph: stacked bar representation of % abundance of isoforms (TPM levels) for peaks during the second subjective dark phase at 20°C (*left*, 40h time-point) and 12 °C (*right*, 44h time-point); total TPM for peaks presented above the stacked bars, abundances (%) denoted within bars or to side of bars. *lower right* Diversity of transcript isoforms at these selected time points is quantified using the Shannon Index. Data for *upper* graphs are normalised mean and ± SEM (*n*=3) of expression levels (TPM; Transcripts Per Million). (b-e) Comparison of 20°C LL and 12°C LL isoform expression profiles for (b) FS (.1); (c) _ID2 (AE); (d) _ID1 (I1R + AE) and; (e) _ID4 (I2R + AE). Data are normalised mean and ± SEM (*n*=3) of expression levels (TPM; Transcripts Per Million).

The cold-switch under diurnal conditions was characterised by a clear flip of two dominant isoforms, FS(.1) and _ID2 (Fig.**1d**). In contrast, the composition of the isoform population generated was more diverse under LL conditions (Fig. **2a**). The isoforms retaining introns 1 and 2 (_ID1 and _ID4, respectively), were more prevalent in the make-up of peaks of the second subjective dark phases for both 20°C LL and 12°C LL (Fig. **1c**, **1d**, Fig. **2a**), compared to the peaks in diurnal conditions (Fig. **1c**). These changes increased the diversity of the *RVE2* transcript population compared to plants held in driven photocycles (Fig. **1d**, Fig. **2a** p < 0.001, Hutcheson’s t test). This suggests that light - or the absence of dark - contributes to *RVE2* splicing fidelity and is a potential influence on *RVE2* oscillatory characteristics.

### Disruption of *RVE2* alternative splicing alters differential gene expression in response to chilling

We wanted to determine the physiological relevance of the *RVE2* cold-switch. To do so, we firstly characterised a Col-0 T-DNA insertional mutant (GABI-Kat 508D10), harbouring the insertion in the AE within the first intron of the *RVE2* gene (Fig. **3a**, Suppl Fig. **S3a**). Homozygous mutants were verified by diagnostic PCR using *RVE2*-specific and T-DNA border primers (Suppl Fig. **S3b** and **d**), and by assessing sensitivity to sulfadiazine (Suppl Fig. **S3c**). qRT-PCR confirmed the absence of the cold-induced induction of FS *RVE2* (Fig. **3b**). Hereafter we refer to the GABI-Kat 508D10 line as *rve2-2* as the second characterised *rve2* allele (Zhang *et al*., 2007).

**Figure 3.**
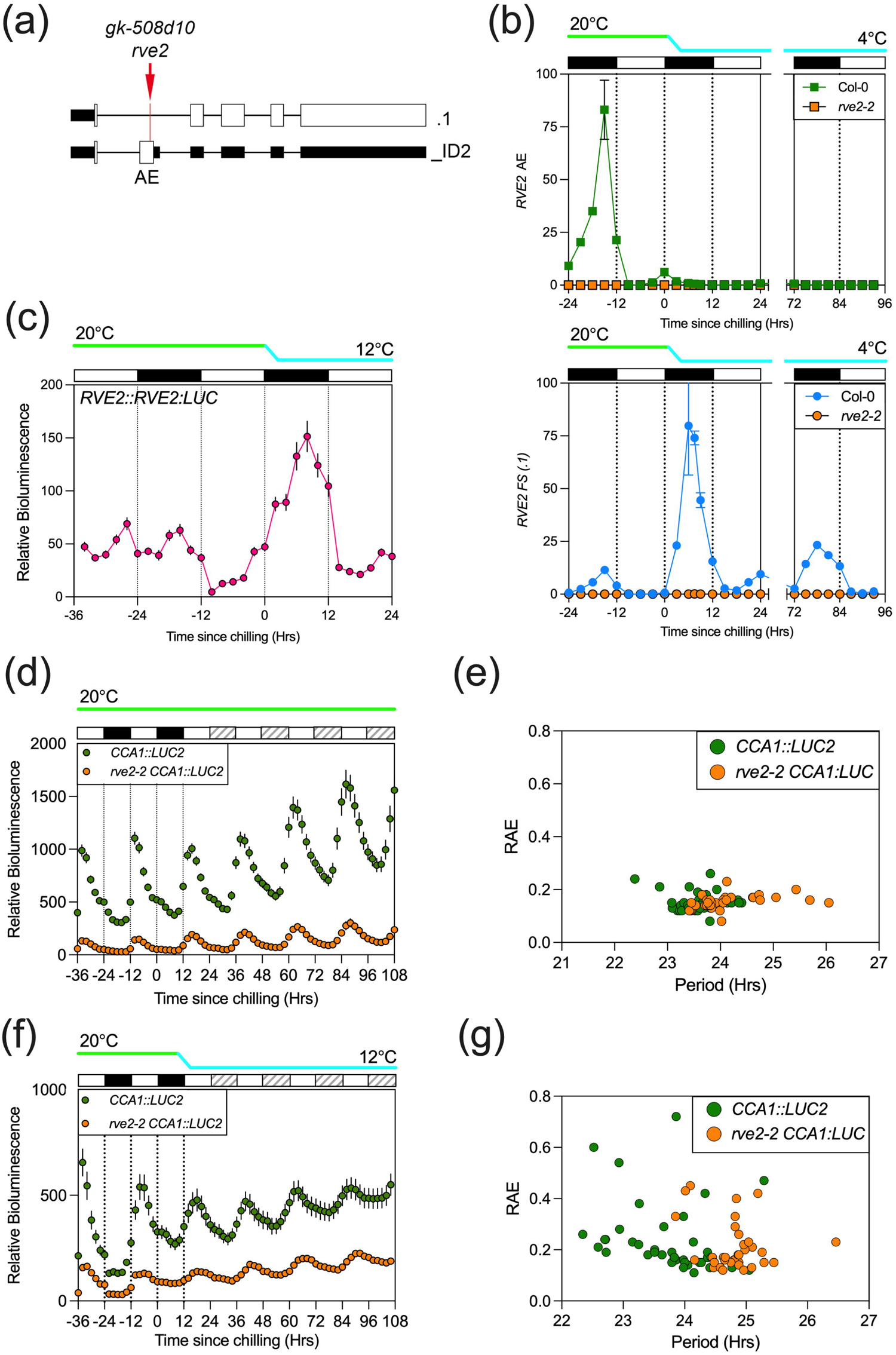
Assessment of circadian phenotypes in *rve2-2* seedlings. (a) T-DNA insertion site of Gabi-Kat line 508D10 within the cold-switch region of *RVE2* intron 1. (b) qRT-PCR to assess accumulation of *RVE2* transcript in wild type and *rve2-2* seedlings. *RVE2* transcripts containing the non-productive Alternate Exon (AE, Fig. **1b**, upper panel) and Fully Spliced (FS, *lower* panel) were estimated using the ΔΔCt method using *IPP2* and *ISU1* as reference transcripts. Samples with the highest expression level were set to 100%. (c) Bioluminescence recorded from seedlings expressing a *RVE2::RVE2:LUC* fusion reporter line before and after chilling. Data is representative of three independent transgenic lines and indicates luciferase activity derived from the fully-spliced RVE2:LUC fusion protein. (d) Patterns of luciferase bioluminescence in wild type and *rve2-2* seedlings in plants expressing a *CCA1::LUC2* reporter held in constant white light at 20°C. (e) Circadian period estimates based on data presented in (d). (f) Patterns of luciferase bioluminescence in wild type and *rve2-2* seedlings in plants expressing a *CCA1::LUC2* reporter held in constant white light following chilling to 12°C at dusk. (g) Circadian period estimates based on data presented in (f). Cooling was initiated at dusk, white and black bars indicate 12h light and dark periods, respectively. Subjective dark periods in constant light are indicated with shaded grey bars.

In order to evaluate the induction of productive *RVE2* splicing we generated a transgenic line expressing a RVE2-LUC translational fusion in the *rve2-2* background (Fig. **3c**). Since chilling to 4°C impairs luciferase activity (Rabha *et al*., 2021), we assessed RVE2:LUC bioluminescence following chilling to 12°C. While quantitative comparisons of luciferase activity at different temperatures are limited by the lability of the enzyme, RVE2-LUC activity increased following chilling, peaking approximately 8h after dusk (Fig. **3c**). These data suggest that FS RVE2 accumulation increases in response to reduced temperatures.

Given the homology of RVE2 to the core circadian clock components CCA1 and LHY (Gray et al., 2017) we next examined whether *rve2-2* had a circadian phenotype using seedlings expressing a *CCA1::LUC2* reporter (Fig. **3**). When held at 20°C under constant white light, *rve2-2* seedlings had a modestly extended circadian period (+0.32 hrs, p <0.01; Fig. **3d** and **3e**). Chilling to 12°C extended circadian period in both wild type and *rve2-2* seedlings, although we observed greater differences between the genotypes at reduced temperatures (+1.1 hrs, p < 0.01, Fig. **3f** and **3g**). These data suggest that RVE2 contributes to the maintenance of circadian rhythms at lower temperatures.

### Transcriptome sequencing reveals that *rve2* plants have altered gene expression in the 12 hours after chilling

In order to assess the role of RVE2 in cold responses we performed RNA-seq on rosettes of 5-week-old Arabidopsis Col-0 and *rve2-2* plants for biological replicates of a diurnal, temperature and time-series experiment (Fig. 4). We adopted the approach of Calixto *et al*., (2018), and described in Tzioutziou et al. (2022), where sampling included plants undergoing a low temperature transition. Rosette tissue samples were harvested at 3h intervals across 24h at 20°C (day 1, ‘ambient’), the following day at 4°C (day 2, ‘transient’), and a third day 3 days later at 4°C (day 5, ‘cool’). We included one additional time-point at the time of peak *RVE2* abundance during day 2 ‘transient’. This was 7.5h after the onset of cooling, corresponding to 1.5h after maximal *RVE2* FS(.1) expression and the peak of *RVE2-LUC* activity (Fig. **1c**, **3c**). The experiment was repeated separately with different batches of Col-0 and *rve2-2* plants on three occasions giving three biological repeats per time point. We quantified transcript abundance in transcripts per million (TPM) using Salmon (Patro et al., 2017) and AtRTD2 as the reference transcriptome (Zhang et al., 2017), allowing us to determine patterns of expression at the individual transcript isoform level for the entire transcriptome (Supplementary Dataset **S4**). RNA-seq reads mapped to the *RVE2* .1 and AE isoform models for Col-0, but not for *rve2-2*, confirming the loss of the cold switch in *rve2-2* (Suppl Fig. **S3e**). Although reads mapped to the other *RVE2* isoform models (Suppl Fig. **S3f-j**), their levels in *rve2-2* were much lower compared to Col-0, except for the _ID1 and _ID4 isoforms (Suppl Fig. **S3f** and **S3h**, respectively), both of which are not predicted to encode functional transcript (Fig. **1b**).

**Figure 4.**
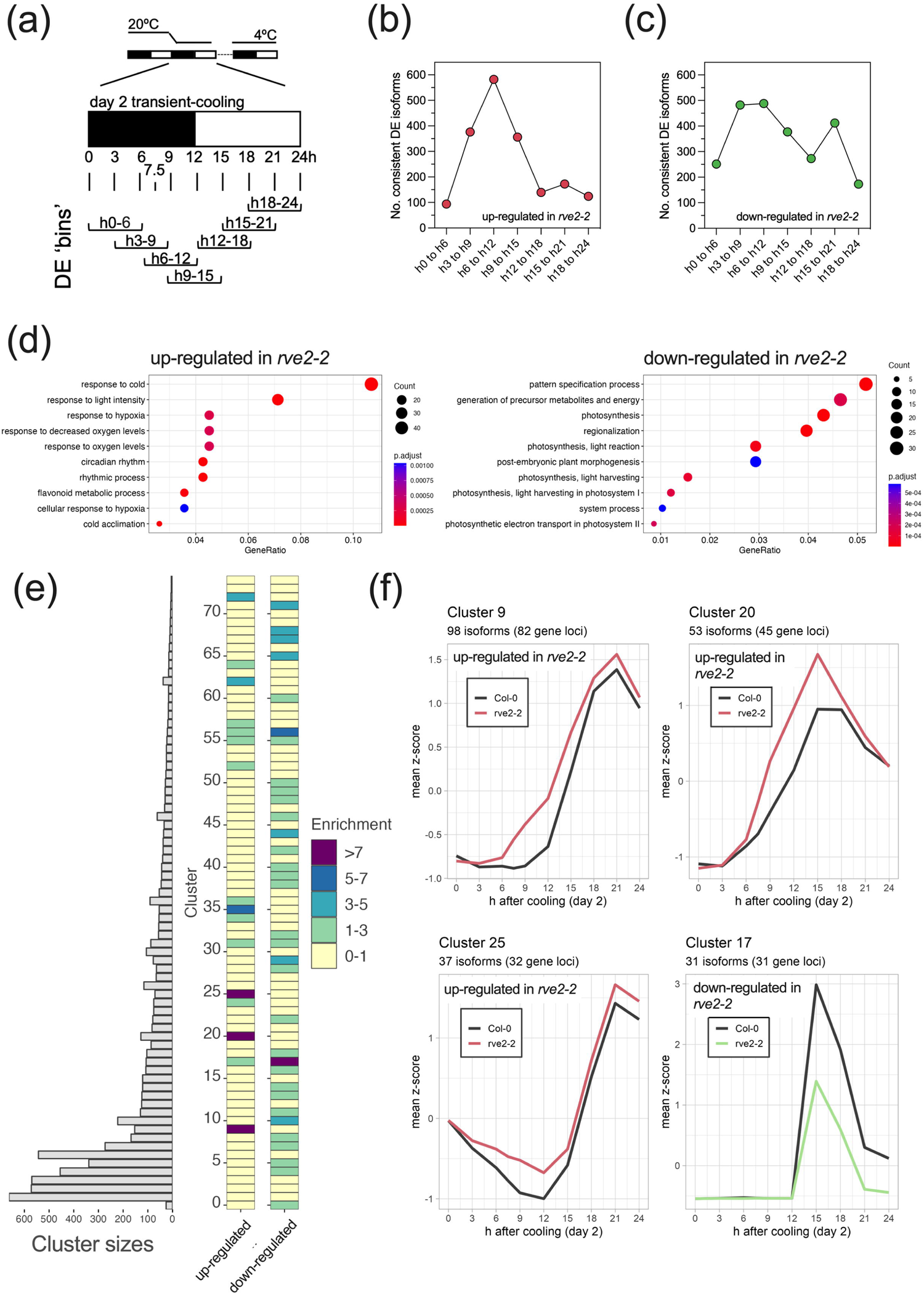
Disruption of the *RVE2* alternative splicing switch alters gene expression following chilling. (a) Overview of categorisation used to identify transcript isoforms consistently Differentially Expressed (DE) in *rve2-2* compared to Col-0 across the transient-cooled day 2 of the temperature and time series RNA-seq experiment. (b) Transcript isoforms consistently up-regulated in *rve2-2* compared to Col-0 in the indicated timeframes. (c) Transcript isoforms consistently down-regulated in *rve2-2* compared to Col-0 in the indicated timeframes. (d) GO term enrichment plots for biological process for the ‘up-regulated in *rve2-2*’ (*left*) and ‘down-regulated in *rve2-2*’ (*right*) DE groups. (e) Alignment and enrichment (Odds ratios) of the grouped h0-24 repressed and activated isoform cohorts with the Calixto *et al*. (2018) TF-cluster groups. (f) Mean z-score of *rve2-2* and Col-0 expression plots for DE isoforms in the denoted enriched cluster groups.

We reasoned that differences in expression between Col-0 and *rve2-2* would be most noticeable at, or soon after, the onset of chilling and so compared transcript accumulation during the 24 hours after temperature reduction. After filtering out low expressed isoforms (discarding isoforms with a sum TPM < 80 across the day 2 timepoints) we identified isoforms showing consistent Differential Expression (DE) for three consecutive timepoints. This approach provided 7 consecutive DE bins across the 24 hour period after chilling, namely h0-6 (i.e. the points at h0, 3 and 6), h3-9, h6-12, h9-15, h12-18, h15-21 and h18-24 (Fig. **4a**). Candidate isoforms were also split into two categories; those that showed consistent DE where *rve2-2* TPM levels were higher than Col-0 (up-regulated) or those that showed *rve2-2* TPM levels lower than Col-0 (down-regulated). We found that the number of consistently up-regulated isoforms increased during the initial phase of cooling (during the first 15 h after chilling) to reach a peak for the h6-12 bin and then decreased (Fig. **4a**). There was a less clear pattern for isoforms down-regulated in *rve2-2*, with an initial peak from h6-12 followed by a second peak in the h15-21 bin (Fig. **4a**). The identity of the down-regulated and up-regulated isoforms for the consistent DE bins are provided in Suppl. Dataset **S5**.

Our DE analysis of the 24 hours after chilling identified 1309 up-regulated and 1720 down-regulated isoforms (Fig. **4a**, Suppl. Dataset **S5**). Two-way ANOVAs were performed on these up-regulated and down-regulated candidate isoform lists to assess the effect of genotype (*rve2-2* and Col-0) and timing on isoform expression levels. Around a third of the DE candidates were significantly mis-regulated (p-values ≤ .05), resulting in groups of 486 and 643 for up-regulated and down-regulated DE isoforms respectively (Suppl. Dataset **S5**). These isoforms are encoded by 435 up-regulated genes and 619 down-regulated genes. Gene Ontology (GO) term enrichment of these gene loci sets show enrichment for Biological Process terms ‘response to cold’, ‘response to light intensity’, ‘circadian rhythm’ and ‘rhythmic process’ for the repressed group and photosynthetic terms for the activated group (Fig. **4b**).

Previously hierarchical clustering of DE genes revealed transient, adaptive, and late expression profiles in response to cold (Calixto et al., 2018). We aligned the ANOVA significant (p ≤ .05), up-regulated and down-regulated gene identities to the Calixto *et al*. cluster groups (clusters 0-74) and performed one-sided Fisher exact tests to establish enrichment (Odds Ratios) of genes within cluster groups (Fig. **4c** and Suppl. Dataset **S5**). Around 68% up-regulated candidates merged with a cluster group (295 out of 435). Some cluster groups were particularly stand out (Odds ratios > 7); clusters 9, 20, and 25 showed high enrichment of consistently up-regulated isoforms (Fig. **4c**). Membership of these three clusters accounted for 54% of the up-regulated group. For the down-regulated candidates around 49% could be assigned a cluster group (302 out of 619) with cluster 17 noticeably enriched (Fig. **4c**), accounting for 10% of the down-regulated group. Fig. **4d** contrasts the *rve2-2* and Col-0 mean z-score expression profiles for the enriched clusters 9, 20, 25 and 17, and suggest that rapid nocturnal repression and subsequent activation of candidate isoforms is the prominent role for RVE2 during a transient-cooled day.

Given the circadian phenotype of *rve2-2* seedlings at reduced temperatures we next assessed the role of RVE2 in regulating circadian genes. Loss of *RVE2* function altered the expression of the _P2 isoform for the *REVEILLE* family member *RVE6* (p= .0018), as well as for the .1 isoform for the core clock gene *PRR5* (p= .019), and the _P1 isoform of *ELF4* (p <.0001) (Suppl. Dataset **S5**). For *RVE6*, expression in *rve2-2* increased to higher levels than Col-0 after 7.5h of cooling and reached a peak 3h in advance of Col-0 at dawn during day 2 ‘transient’ (Fig. **5a**). For Col-0 expression of the *PRR5* .1 isoform is unresponsive to the onset of cooling in the dark. However, in *rve2-2* two peaks of expression were evident; expression increased after the onset of cooling to give an additional peak 6h after cooling then continuing to the daytime later the following day (Fig. **5b**).

**Figure 5.**
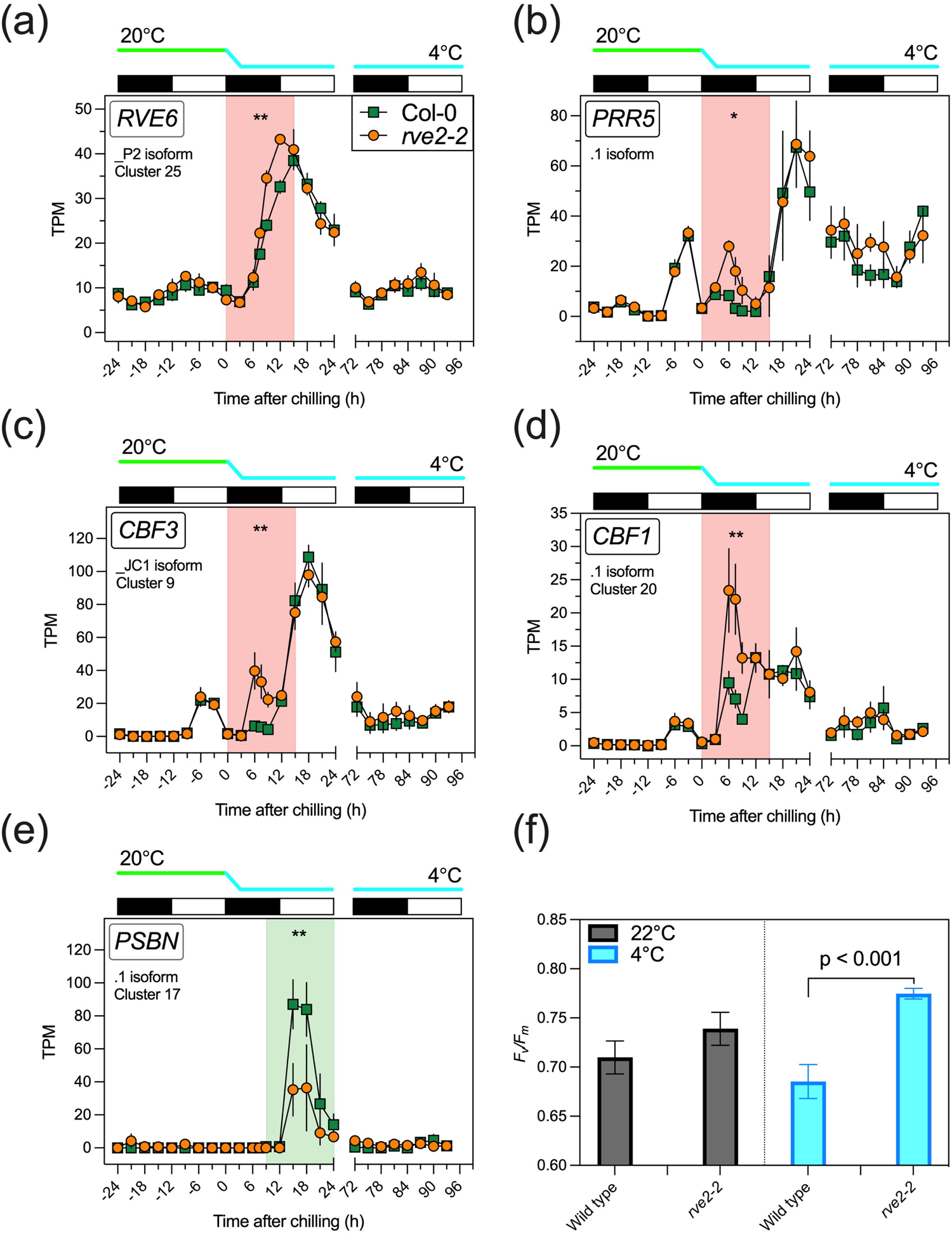
Altered expression of candidate transcripts in *rve2-2* during the onset of cooling. Temperature and time-series transcript expression plots for a selection of candidate isoforms displaying altered expression in *rve2-2* compared to Col-0. Two-way ANOVA p-value summary statistic for either ‘up-regulated in *rve2-2*’ or ‘down-regulated in *rve2-2*’ (red or green shaded regions, respectively), for (a) the _P2 isoform of *REVEILLE 6* (*RVE6*), (b) the .1 isoform of *PSEUDO-RESPONSE REGULATOR 5 (PRR5)* (c) the _JC1 isoform of *C-REPEAT BINDING FACTOR 3* (*CBF3*), d) the .1 isoform of *C-REPEAT BINDING FACTOR 1* (*CBF1*) and (e) the .1 isoform of *PHOTOSYSTEM II REACTION CENTER PROTEIN N* (*PSBN*). (f) Maximum quantum efficiency of PSII photochemistry of Col-0 and *rve2-2* seedlings before and after cool night chilling. For panels (a - e), data represents TPM mean and ± S.E.M (*n*=3). Alternating black-white rectangles denote 12 h dark-light phases, respectively. Cluster designations correspond with Figure 4. Data is presented relative to the onset of chilling at dusk (ZT12).

Beyond circadian genes, altered expression was found for the *C-REPEAT BINDING FACTOR 3* (*CBF3*, _JC1 isoform, Fig **5c**, p= .0074). CBF3, also known as DREB1A, is a member of the ERF/AP2 transcription factor family that includes CBF1/DREB1B and CBF2/DREB1C, that bind the dehydration-responsive *cis*-element (DRE) in the promoters of many downstream cold-inducible genes. Since all three *CBF* genes are rapidly and significantly induced by cold stress, their induction is generally considered to be the first switch in the cold-responsive expression of numerous genes (Kidokoro et al., 2020, Yamaguchi-Shinozaki and Shinozaki, 1994). Altered expression between *rve2-2* and Col-0 was established for both *CBF1* and *CBF2* (.1 isoform p= .003 and _P1 isoform p= .0083, respectively). For all three *CBF*s, transcript levels were higher in *rve2-2* compared to Col-0 after 6h of cooling and remained elevated until dawn of day 2 ‘transient’. These differences seemed particularly focussed to this part of the cooling night, other sections of the expression profiles were virtually indistinguishable between *rve2-2* and Col-0 (*CBF3*, Fig **5c**; *CBF1,* Fig **5d** and CBF2, Suppl. Dataset **S5**).

We found that all the 31 Cluster 17 members are chloroplast encoded and are down-regulated in *rve2-2* compared to Col-0 across the light phase of day 2 ‘transient’ (cluster 17, Fig. **4d**). For example, we saw distinct *rve2-2* and Col-0 expression profiles for *PHOTOSYSTEM II REACTION CENTER PROTEIN N* (*PSBN,* .1 isoform, Fig. **5e**, p= .01). The expression profiles for all other chloroplast-encoded candidate gene were broadly similar with a pulse of expression after dawn in the light phase of day 2 ‘transient’, with expression maxima 3h after dawn. We next assessed the physiological effect of these changes in chloroplast-related gene expression (Figure **5f**). We expected that the differences in gene expression observed would culminate in altered photosynthetic performance, and so measured the maximum quantum efficiency of PSII photochemistry (*F_v_/F_m_*) in plants either maintained at 22°C or chilled overnight at 4°C. We were interested to observe that *rve2-2* photosynthetic capacity was significantly higher than wild-type plants following chilling (p<.001, Student’s t test; Figure **5f**). Such data suggest that RVE2 serves to limit photosynthetic performance at reduced temperatures.

### Analysis of possible components required for *RVE2* alternative splicing

AS has been linked to circadian clock control in plants, in part due to the observation that mutants defective in a variety of splicing factors (SFs) and spliceosome components show defects in circadian behaviours [reviewed in (Mateos *et al*., 2018)]. We were interested whether specific parts of the RNA splicing machinery contributed to the integration of temperature information with the alternative splicing observed in *RVE2* following chilling. These components include Splicing Factors/RNA-Binding Proteins (SF/RBPs), for example those with altered cold sensitivity or tolerance when mis-expressed (Reddy *et al*., 2013; Staiger & Brown, 2013; Laloum *et al*., 2018) and reversible phosphorylation of SFs by environmentally-regulated signalling pathways (e.g. mitogen-activated protein kinases, MAPKs) (Teige *et al*., 2004; Wurzinger *et al*., 2011; Knight & Knight, 2012). We therefore tested *RVE2* cold-switching strength in mutants of the *POLYPYRIMIDINE TRACT BINDING* (*PTB*) family, heterogenous ribonucleoprotein (hNRP) SFs previously shown to influence the response of the *LHY* isoform switch to temperature (James *et al*., 2018), as well as in plants lacking *MITOGEN ACTIVATED PROTEIN KINASE KINASE1 (MKK1)* and *MKK2* since *mkk2* plants demonstrate reduced ability to cold acclimate (Teige *et al*., 2004).

### Modulation of the *RVE2* cold-switch strength in hnRNP splicing factor mutants

We have previously shown that splicing and expression of polypyrimidine tract-binding proteins (PTBs) – a family of heterogeneous nuclear RNP (hnRNP) SF proteins - respond rapidly and sensitively to temperature changes to contribute to the splicing of the 5′UTR of *LHY* (James *et al*., 2018). PTBs bind to the pyrimidine-rich regions of introns often, but not exclusively, upstream of 3′ splice sites. Traditionally hnRNP SFs have been regarded as repressors of splicing but it is now evident that the mode of action of SFs is often context-dependent and typically involves the interaction of multiple components of the splicing machinery and additional regulatory factors(Wachter *et al*., 2012). We noted that, like *LHY*, the 5’ region of the *RVE2* gene sequence was enriched for pyrimidine bases, for example three polypyrimidine (pY) tracts flanking and immediately adjacent to the predicted transcriptional start site [Fig. **6a**; (Hieno *et al*., 2014)]. Both *PTB1* (At3g01150) and *PTB2* (At5g53180) undergo chilling-induced isoform switching such that the levels of the protein coding isoforms rise rapidly after the onset of cooling in the dark, although this transition is slower than observed in *RVE2* [(James *et al*., 2018) Fig. **6b**]. Re-analysis of previous datasets revealed that three hours after chilling, *PTB1*(.1) levels were higher compared to the equivalent time-point at 20°C (Student’s t-test; p = .0484), whereas *PTB2* (_P1) levels were not different until 9h after cooling (Student’s t-test; p = .0187) [Fig. **6c**; (Calixto *et al*., 2018)]. Such data demonstrate that *PTB* gene expression is responsive to nocturnal chilling, albeit at a slower rate than *RVE2*.

**Figure 6.**
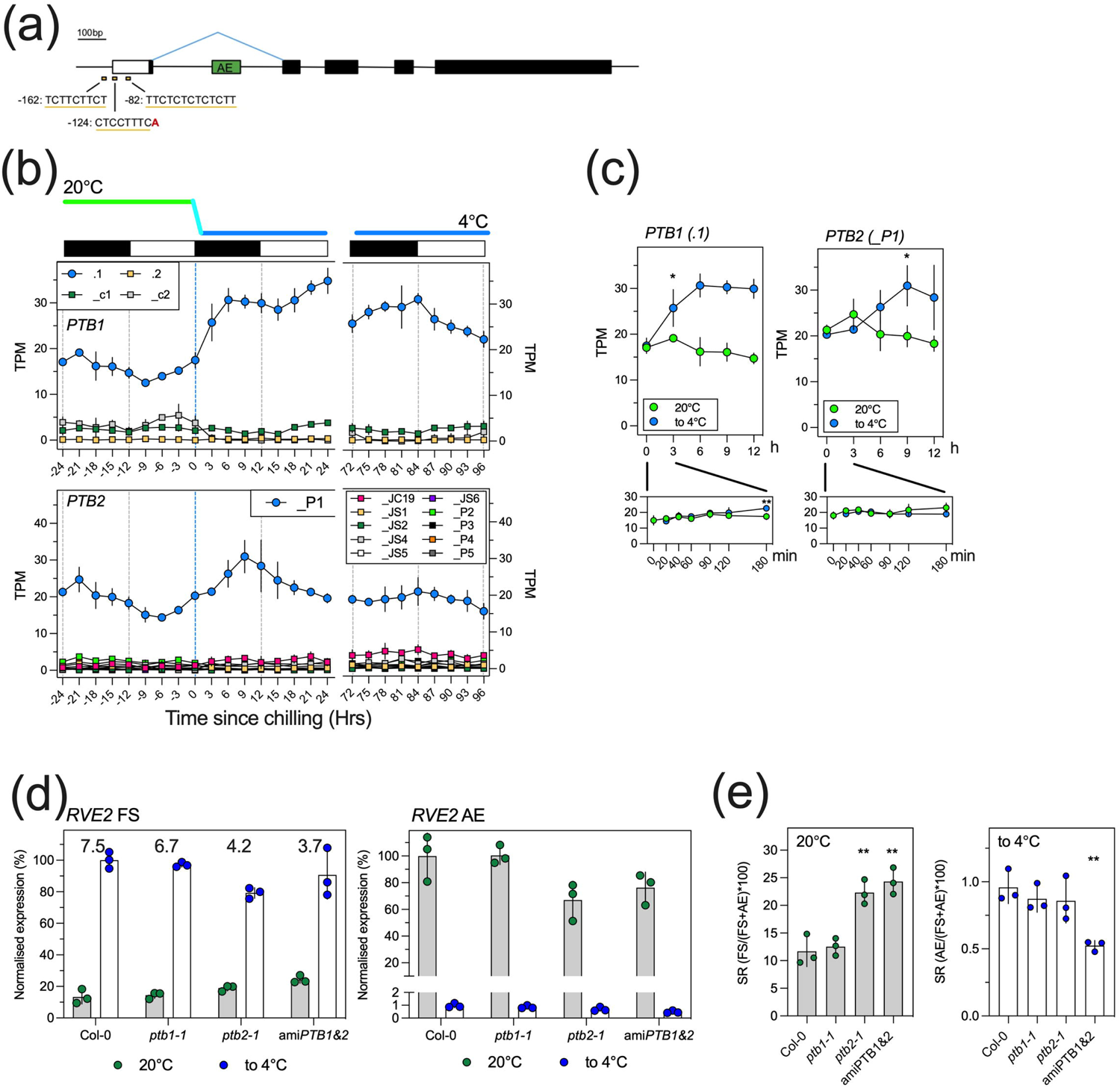
Modulation of the *RVE2* cold-switch strength in hnRNP splicing factor mutants. (a) Polypyrimidine tract (pY) features of the *RVE2* promoter landscape (redrawn from ppdb output (http://ppdb.agr.gifu-u.ac.jp). pY sequence and coordinates (relative to the translational start codon) in the 5’UTR of the *RVE2* locus, putative transcriptional start site denoted as red text ‘A’. Transcript map: black box – coding sequences; white box – 5’ UTR; green box – alternative exon (AE). (b) Temperature and diel time series RNA-seq profile for *PTB1* (*top*, At3g01150) and *PTB2* (*bottom*, At5g53180), data of Calixto et al., 2018. Mean and ± SEM (*n*=3) of normalised expression levels (TPM; Transcripts Per Million) for the denoted transcript isoforms, cooling from 24h (dashed blue vertical line), diurnal 12h dark/12h light conditions as black/white rectangles, respectively. (c) RNA-seq expression levels of *PTB1* (*left*, ; .1) and *right*; *PTB2 (_P1)*, data of Calixto et al., 2018 for top graphs (12h dark-phased segments of the 20 and 20 to 4°C days), and IE RNA-seq dataset (Supplementary Dataset **S2**) for bottom graphs (initial 3h of dark-phased segments of the 20 and 20 to 4°C days). Data is mean and ± SEM (*n*=4) of normalised expression levels (TPM; Transcripts Per Million). Statistical summaries (* = p ≤ 0.05, ** = p ≤ 0.01) for paired two-sample t-tests between temperature treatments. (d) *RVE2* expression levels for FS(.1) – *left* and AE (_ID2 and _JC4) – *right* for the denoted genotypes at either 20°C or 20 to 4°C (‘to 4°C’). Data is mean ±SEM (*n*=3) of normalised expression levels. Transcript levels were estimated using the Δ Δ Ct method and real time RT-qPCR using *IPP2* and *ISU1* as reference genes, *RVE2*-FS primers and *RVE2*-AE primers (Suppl Fig. **S1c**, Supplementary **Table S1**). FS(.1) levels (*left*) are relative to Col-0 at 6h after dusk ‘to 4°C’ (calibrated as 100%) and AE (_ID2 and _JC4) levels (*right*) are relative to Col-0 at 9h after dusk at 20°C (calibrated as 100%). (e) *RVE2* splice ratios (SRs) for the denoted genotypes; *left* - FS(.1) as a fraction of total (FS(.1) +AE) levels for the 20°C data (9h after dusk) and *right* - AE as a fraction of total levels for the cooling (‘to 4°C’) data (6h after dusk). Statistical summaries (** = p ≤ 0.01) for unpaired two-sample t-tests comparing SRs between *ptb* mutants and Col-0.

We next used RT-qPCR to measure levels of the *RVE2* FS(.1) and AE isoforms in single locus mutations *ptb1-1*, *ptb2-1* and an artificial microRNA (ami) knockdown of *PTB1* and *PTB2*, in the Col-0 background, where the knockdown reduces both mRNAs to around 40% of wild-type *RVE2* transcript levels [Fig. **6d**, (Ruhl *et al*., 2012)]. We selected two time-points for comparison, the peak of *RVE2* AE accumulation (9h after dusk at 20°C) and the peak of *RVE2* FS(.1) accumulation (6h after dusk during cooling). In general, we found that *ptb2-1* and the amiPTB line showed the largest differences compared to Col-0, with *RVE2* FS(.1) levels increased when held at 20°C in comparison to wild type plants (Fig. **6e**, p < 0.01). Significantly, a concomitant decrease in *RVE2* AE was observed in *ptb2-1* and *amiPTB* lines (Fig. **6e**, p<0.01). The increased accumulation of *RVE2* FS at 20°C in these lines altered the magnitude of differential splicing following chilling, which we quantified by comparing *RVE2* Splice Ratios (SRs) at the time of peak accumulation for each isoform (Fig. **6c**). Student’s t-tests revealed a significant difference in SR at 20°C between *ptb2-1* and Col-0 (p = .0066) and between amiPTB1&2 and Col-0 (p = .0040). There was also a significant difference in the SR in the cooling data between amiPTB1&2 and Col-0 (p = .0041). Our data suggest that PTBs contribute to regulating the *RVE2* isoform switch but are not necessary for cold switching.

### *RVE2* cold-switch strength analysis in mitogen-activated protein kinase mutants

As an alternative to the contribution of PTBs, we also speculated that the rapid onset of the *RVE2* cold switch could reflect phosphorylation of unidentified SF(s) via an environmentally regulated MAPK signalling pathway. The protein-coding isoforms of both *MKK1* (At4g26070) and *MKK2* (At4g29810) peak shortly after dusk in unchilled conditions, and demonstrate differing responses to chilling (Calixto et al., 2018; Fig. **7a** and **7b**). Levels of the *MKK1* _P2 isoform are elevated 3 hours after the initiation of chilling (Student’s t-test, p = .0008), whereas *MKK2* _P1 transcript accumulation is maintained after cooling, resulting in significantly elevated transcript compared to the mock-treated control after 6 hours (Student’s t-test, p < .0001; Fig. **7b**).

**Figure 7.**
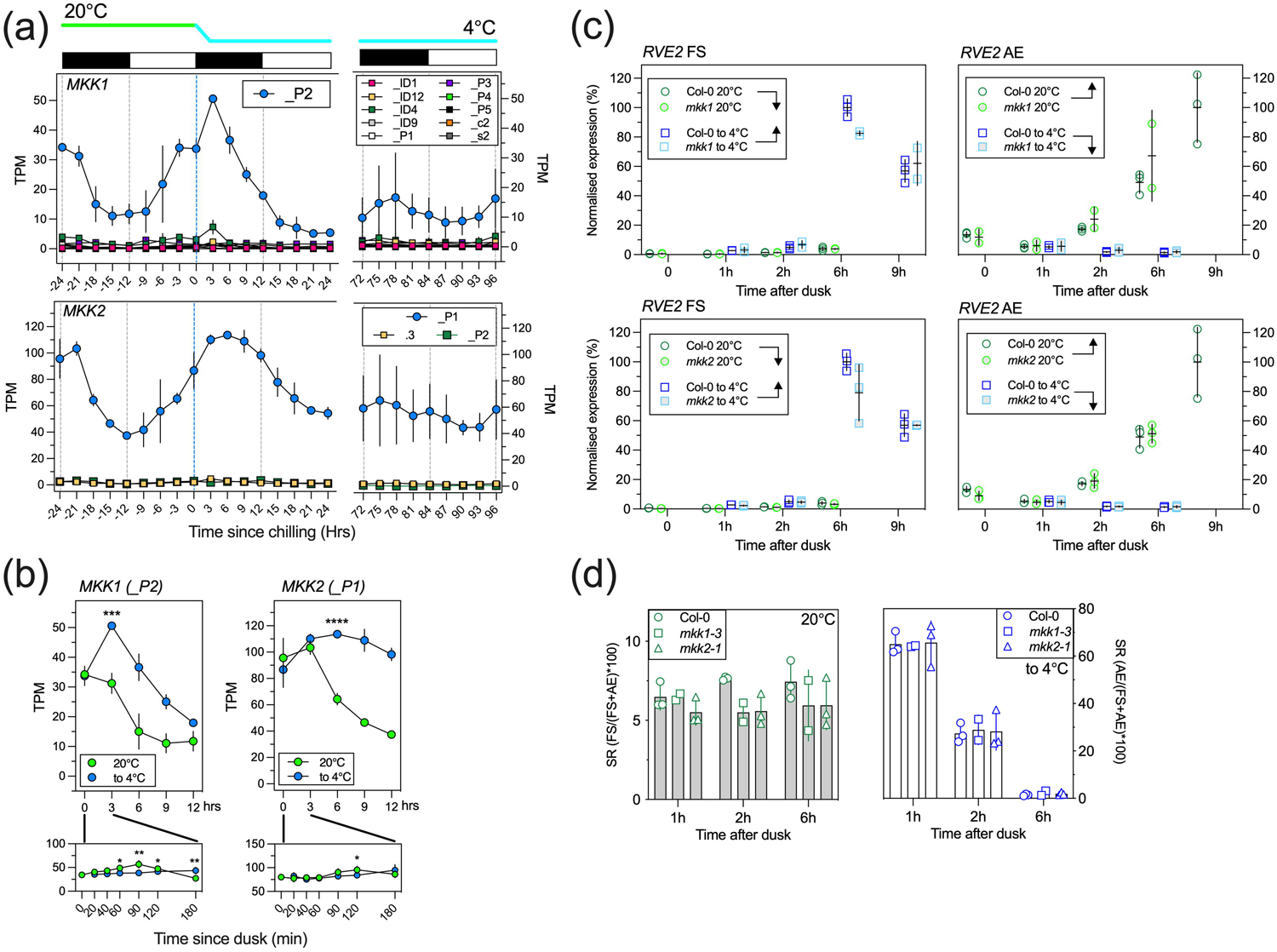
*RVE2* cold-switch strength analysis in mitogen-activated protein kinase (*MAPK*) mutants. (a) Temperature and diel time series RNA-seq profile for *MKK1* (*top*, At4g26070) and *MKK2* (*bottom*, At4g29810), data of Calixto et al., 2018 Mean and ± SEM (*n*=3) of normalised expression levels (TPM; Transcripts Per Million) for the denoted transcript isoforms, cooling from 24h (dashed blue vertical line), diurnal 12h dark/12h light conditions as black/white rectangles, respectively. RNA-seq expression levels of (b) left; *MKK1* (_P2) and right; *MKK2* (_P1), data of Calixto et al., 2018 for top graphs (12h dark-phased segments of the 20 and 20 to 4°C days), and IE RNA-seq dataset (Supplementary Dataset **S2**) for bottom graphs (initial 3h of dark-phased segments of the 20 and 20 to 4°C days). Data is mean and ± SEM (*n*=4) of normalised expression levels (TPM; Transcripts Per Million). Statistical summaries (* = p ≤ 0.05, ** = p ≤ 0.01, *** = p ≤ 0.001, **** = p ≤ 0.0001) for paired two-sample t-tests between temperature treatments. (c) Expression levels for *RVE2* FS(.1) – *left* and *RVE2* AE (_ID2 and _JC4) – *right* for the denoted genotypes at either 20°C or 20 to 4°C (‘to 4°C’) in the *mkk1* background – *top*, or the *mkk2* background - *bottom*. Data is mean ±SD (*n*=2-3) of normalised expression levels. Transcript levels were estimated using the ΔΔCt method and real time RT-qPCR using *IPP2* and *ISU1* as reference genes, *RVE2*-FS primers and *RVE2*-AE primers (Suppl Fig. **S1c**, Supplementary **Table S1**). FS(.1) levels (*left*) are relative to Col-0 at 6h after dusk ‘to 4°C’ (calibrated as 100%) and AE (_ID2 and _JC4) levels (*right*) are relative to Col-0 at 9h after dusk at 20°C (calibrated as 100%). Arrows in the legend denote expected expression levels (low; downward arrow; high; upward arrow). (d) *RVE2* splice ratios (SRs) for the denoted genotypes and time points; *left* - FS(.1) as a fraction of total (FS(.1) +AE) levels for the 20°C data and *right* - AE as a fraction of total levels for the cooling (‘to 4°C’) data.

MKK2 is a member of the MEKK1-MKK1/2-MPK4 signaling cascade that positively affects plants cold response (Zhao *et al*., 2017), and although MKK1 and MKK2 are generally thought to have redundant functions (Gao *et al*., 2008), cold-induced activation of MPK4 was shown to be completely blocked in the *mkk2* single mutant, but not affected in the *mkk1* mutant, suggesting that MKK1 and MKK2 have different functions in cold signaling (Zhao *et al*., 2017). Plants lacking MKK2 display reduced ability to cold acclimate and overexpression of MKK2 leads to the induction of many cold genes in the absence of cold, consistent with MKK2 being a positive regulator of cold gene expression (Teige *et al*., 2004; Knight & Knight, 2012; Zhao *et al*., 2017). Using qRT-PCR we assessed *RVE2* isoform levels in *mkk1-3* and *mkk2-1* mutants [Fig. **7c**; (Teige *et al*., 2004; Qiu *et al*., 2008)] and determined the SRs from these data (Fig. **7d**). This analysis did not reveal any difference in either *RVE2* FS or *RVE2* AE splicing and so we conclude that cold-switching in *RVE2* appears to be insensitive to mutations in the MAP kinases investigated.

## Discussion

### Deep Sequencing Reveals a Novel, Chilling-Induced Alternative Splicing Switch

REVEILLE2 is a member of a wider group of single Myb-like domain transcription factors, and while most of the family have ascribed functional roles in the circadian clock - LHY and CCA1 most prominently – a role for RVE2 has remained largely enigmatic. We mined the diel temperature and time series dataset of Calixto et al., 2018 for those genes displaying the most prominent combined DE and DAS in plants undergoing cooling. *RVE2* clearly stands out with respect to these features. Other genes are expressed to higher levels (*y*-axis of Fig. **1a**), for example members of the Rubisco small subunit multigene family, but unlike *RVE2*, these genes display little or no AS (*x*-axis of Fig. **1a**). Likewise, those gene isoforms that are higher ranked than *RVE2* FS(.1) and _ID2 isoforms for positive and negative ΔPS show very low expression levels (less than 2.5 maxTPM). Our RNA-seq was performed on total RNA from Arabidopsis rosettes and we recognise that cell, tissue and development-specific expression levels may be masked or absent, meaning that comparing gene maxTPM levels can be potentially misleading. However, the speed of induction of the *RVE2* cold switch (Fig. **1e**) and the near complete transition from non-functional to functional transcript isoforms with cooling (Fig. **1d**) validate our decision in pursuing *RVE2* as a new component of plants’ cold response.

Although some aspects of RVE2 expression and AS have previously been described (Filichkin & Mockler, 2012; Filichkin *et al*., 2015), the isoform switching between non-translatable and translatable transcript messages with cooling has until now remained hidden. We suggest there are two main reasons for this. Firstly, we show that only a small proportion of *RVE2* gene expression is productively spliced at ambient temperature and therefore studies characterizing *RVE2* (including phenotypic studies of adult *rve2* plants) at standard ambient temperatures are rendered largely redundant. Secondly, the detection of the *RVE2* cold switch was only made possible by the transcript-specific RNA-seq analysis – microarray or gene-level RNA-seq analyses would not reveal this AS temperature switch. In addition, transcript-specific RNA-seq analyses relies upon employment of a high quality, non-redundant reference transcript datasets that distinguish diverse transcript assemblies. Since the TAIR10 reference transcriptome contains only a single *RVE2* splicing isoform previous studies examining temperature-dependent alternative splicing could not distinguish the *RVE2* AS temperature switch (e.g. Romanowski *et al*. 2020 and Bonnot *et al*. 2021). In contrast we employed an alternative transcriptome reference that recognises seven *RVE2* transcript isoforms (AtRTD2; Zhang *et al*., 2017). Most of the differences between these isoforms are due to splice site choice in the first intron (Fig. **1b**). Two of the isoforms, _ID2 (containing an AE in intron 1, introducing a premature termination codon), and the protein coding FS (.1), make up 78-92% of *RVE2* expression in the “off-on” states respectively and are therefore primarily accountable for the RVE2 temperature switch. Work examining plants over-expressing the *RVE2* cDNA can be re-interpreted in the context of the cold-induced isoform switch described here (Zhang *et al*., 2007; Guan *et al*. 2013).

The FS(.1) isoform peaks in the middle of the night after cooling, some 3h earlier than the much smaller peak at 20°C. This flexibility in the accumulation of *RVE2* transcripts may ensure that internal cellular events are timed appropriately. In terms of those circadian components phased to around dawn, *CCA1* and *LHY* for example, peak adjustment occurs because the oscillator entrains to the new time of the dawn signal through phase adjustments each day. In this scenario light-regulated transcription factors, such as phytochromes and cryptochromes, regulate the dawn phased components (Somers *et al*., 1998; Martinez-Garcia *et al*., 2000). For other clock genes, the mid-afternoon phased component *PRR7* and dusk phased *TOC1*, phase resetting appears to be regulated by sugar sensing (Haydon *et al*., 2013; Frank *et al*., 2018) and ZTL-mediated degradation (Mas *et al*., 2003), respectively. Growing evidence also points to temperature-dependent splicing and nonsense-mediated decay transducing temperature signals to alter the phase of the oscillator (Wang *et al*., 2022). Productive *RVE2* switches on and advances phase almost simultaneously with cooling (Fig. **1**, Suppl. Fig. **S1**). The immediacy of this response strongly suggests a mechanism centred on thermosensitive splicing. The cooling associated phase advance to the mid-point of the nocturnal phase is maintained with acclimation to 4°C (Fig. **1c**), and while temperature as an entraining signal is notoriously difficult to mechanistically isolate (since temperature affects nearly all biological processes; Wang *et al*., 2022), cool nights may be particularly important to adjust the pace of the clock via *RVE2* thermo-splicing. RVE2 has not previously been regarded as contributing to circadian clock function, possibly due to the classical (e.g. steady state ambient temperature) ways in which clock functions are typically assigned. However, using a clock promoter fused to luciferase expression (*CCA1::LUC2*) we established a clock phenotype (Fig. **3**) showing that RVE2 contributes to circadian timing at reduced temperatures. Our data therefore establish the potential for assigning a more prominent role for RVE2 in clock function that merits further development, particularly the possible role of RVE2 thermo-splicing for entraining the clock during cool nights.

Potential RVE2 targets were identified using RNA-seq and temperature and time series experiments; here we compared plants devoid of the *RVE2* cold-switch (*rve2-2*) with wild-type Col-0 plants (Figs. **4** and **5** and **Table 1**). Derepressed expression profiles were a feature of a large proportion of candidate genes, suggesting that RVE2 primarily functions as a repressor of target gene expression, the exception being the chloroplast encoded genes previously mentioned. Altered profiles were mostly focused to the mid-nocturnal phase of the cooled night implying that RVE2 direct targets are repressed with a subsequent, indirect induction of gene expression (Fig. **4a**). Later in the timecourse, our RNA-seq data show altered expression in *rve2-2* for a set of chloroplast-encoded genes (**Table 1**) as well as increased photosynthetic capacity after cooling (Fig. **5f**). This points to RVE2 playing a role - likely indirectly - in priming photosynthesis capacity with the prospect of imminent dawn after a cooled night.

Our data therefore provide a link between clock mediated integration of rapid, and possibly fluctuating, changes in nocturnal temperature with coordinating an appropriate cold response. Presumably plants need to gauge their energy gathering potential at the break of day – the temperature information transduced by RVE2 in the hours before this may play a part in this ‘decision’, although in the case of plants adapted to the high Arctic photosynthesis at low temperature during summer months is rather more predictable. Arctic plants show optimum photosynthesis rates at a lower temperature than other plants (Chapin, 1983) and in this regard, it is particularly intriguing that a recent study comparing the cold response transcriptomes of Arctic plants found *RVE2* as a shared differentially expressed gene (Birkeland *et al*., 2022).

### Constant Light Also Contributes to *RVE2* Splicing Outcomes

Reductionist approaches using structured environmental conditions in laboratory settings, such as square-wave light/dark cycles and constant light/dark or temperature, are routinely used for investigating the architecture and function of the circadian oscillator, even if plants in nature are unlikely to experience these conditions (Panter *et al*., 2019). We tracked *RVE2* transcript isoform oscillations for ‘free-run’ conditions of constant cool temperature and constant light. In general, *RVE2* transcript isoforms continued to oscillate over 56h of constant conditions (Fig. **2**), implying that these rhythms can persist in the absence of timing cues. The isoform switch was still apparent in plants held in constant light after chilling but the FS(.1) isoform rhythm was severely damped, with a higher proportion of non-functional isoforms making up the peak of the rhythm (Fig. **2**). Light has been shown to regulate AS in plants; light-exposed plants show faster gene transcription compared to those in the dark, and since transcription and splicing are mechanistically coupled, light:dark context serves as a control for alternative mRNA splicing decisions (Petrillo *et al*., 2014; Godoy Herz *et al*., 2019). Constant light ‘free-run’ conditions at lower temperatures are analogous to continuous daylight in summer at Arctic and Antarctic latitudes, and it is therefore conceivable that in nature a light:dark coincidence mechanism provides a layer of *RVE2* cool-switch regulation.

Rapid, discontinuous entrainment via coincidental cool, nocturnal conditions could conceivably regulate seasonal expression of the *RVE2* functional isoform and merits further consideration. For example, due to seasonal changes the time of the day that *RVE2* peak expression occurs during cold nights would be required to shift over the course of the year. For Glasgow and Dundee (United Kingdom, Latitude 56° N), not only does the time of dusk change by about 6 h over the year but also the length of the night, such that at the summer solstice the Sun barely dips below the horizon. Changes in photoperiod also adjust the time of peak expression of the components of the clock with oscillator genes tending to peak later in long daylengths (Webb *et al*., 2019). Warmer nights associate with, but are not limited to, the longer days of spring and summer and it will be interesting to learn if the interplay between temperature, daylength, light intensity and light quality influences *RVE2* FS(.1) expression, entrainment and cold response gating.

### The Cold Sensing Mechanism Underlying *RVE2* Alternative Splicing Remains Unknown

We imagine that the factor(s) integrating temperature information with dynamic alterations in *RVE2* splicing outcomes are potentially wide and varied. Our basis for pursuing PTB splicing factors as potential regulators of the RVE2 cold-switch was based on our previous work on the ‘on:off’ thermo-splicing switch at the 5’UTR region of *LHY*. Like *LHY*, pY sequence patches – potential binding sites for polypyrimidine binding tract proteins – are found in the vicinity of the transcriptional start site. We found that the cold-switch was weaker, but not abolished, in an amiRNA knock-down line for *PTB1* and *PTB2* compared to Col-0 (Fig. **6**). This was similar to our observations with *ptb* mutants and LHY thermo-splicing (James *et al*., 2018). PTBs therefore appear to contribute to, but are not sufficient for, modulating the *RVE2* cold-switch. This does not rule out roles for other SFs in regulating the *RVE2* cold-switch, indeed *RVE2* has been shown to be mis-spliced during cold stress in a mutant line for a DEAD box RNA helicase, REGULATOR OF CBF GENE EXPRESSION 1 (Guan *et al*., 2013). Similarly, we pursued the idea that MAP kinases could potentially regulate SFs acting on the *RVE2* cold-switch on the basis that changes in phosphorylation and intracellular Ca^2+^ is at least one way in which temperature information influences the balance between functional and non-functional SF isoforms (Stamm, 2002; Stamm, 2008; Xie, 2008). For example, a mammalian body temperature-sensitive thermometer-like kinase has been identified that alters SF protein phosphorylation, thereby globally controlling AS changes (Preussner *et al*., 2017). However, the MAP kinases we selected to examine appeared to have little effect on *RVE2* cold-switch dynamics (Fig. **7**).

For plants, timing is everything. Flexibility in timing is also crucial for the ability of plants to adapt to new environments and changing climates (Fournier-Level *et al*., 2011). The novel *RVE2* ‘cold-switch’ therefore offers the opportunity to understand how plants perceive and integrate overnight chilling signals.

## Data Availability

The data that support the findings of this study will be openly available in the NCBI Sequence Read Archive (SRA) at https://www.ncbi.nlm.nih.gov/sra, reference numbers PRJNA909847 and PRJNA909639.

## Acknowledgements

The authors thank NASC (Scholl, May, & Ware, 2000) for the provision of seed. This work was supported by UKRI/BBSRC (grants BB/P006868/1, BB/S005404/1, BB/P009751/1, BB/S020160/1) and the Scottish Government Rural and Environment Science and Analytical Services division (RESAS; grant number JHI-B1-2).

## Conflict of Interest

The authors are unaware of any conflict of interest.

## Author Contributions

ABJ, JWSB, RZ, HGN, and MAJ planned and designed the research. ABJ, CS, JL, EMA, NT, MAJ performed experiments, ABJ, CS, WG, RZ, MAJ analysed data. ABJ, HGN, MAJ wrote the manuscript with help from all authors.

---

**Fig. S1. The splicing driven *RVE2* cold switch**. (a) Simplified form of main Fig **1c** clearly showing the isoform switch of the two main *RVE2* isoforms, FS(.) and _ID2. Data of Calixto et al., 2018; mean and ± SEM (*n*=3) of normalised expression levels (TPM; Transcripts Per Million) for the denoted transcript isoforms, cooling from 24h (dashed blue vertical line), diurnal 12h dark/12h light conditions as black/white rectangles, respectively. (b) The speed of changes in expression/AS were examined for a high-resolution RNA-seq time-series across the first 3h of cooling (Supplementary Dataset **S2**). Plotting the time-point at which genes were first significantly DE or DAS (each gene is represented only once) showed no significant DE genes at 20 min while two genes were significantly DAS at 20 min. One of two DAS genes with the most rapid significant AS was *RVE2*. (c) Graphical representation of predicted annealing locations of RT-qPCR primers (Supplementary **Table S1**) amplifying isoform specific *RVE2* amplicons. Primers spanned introns (dotted lines); forward (fwd), and reverse (rev) orientation denoted, primer sequence 5’ to 3’. The FS(.1) primer combination is predicted to amplify FS(.1) specifically, whereas the AE primer combination is predicted to amplify both the _ID2 and _JC4 isoforms. (d) Expression profile of the two principal *RVE2* isoforms (FS(.1), *left* and AE (_ID2 and _JC4), *right*) using the RT-qPCR primers described in the middle panel. Time points spanned 12h of dusk (at either 20°C or cooling onset, 20 to 4°C) except for closely spaced time points during the first 3h of dusk (grey shaded area expanding to upper graphs). The DDCt method was used to calibrate normalised Ct values (using the average of Cts for IPP2 and ISU1 housekeeping genes) to the time-point demonstrating maximal expression (6h at 20 to 4°C for FS(.1), *left*; and 9h at 20°C for AE, *right*). Data is mean ±SEM (*n*=3) of normalised expression levels. Statistical summaries (*** = p ≤ 0.001, **** = p ≤ 0.0001) for paired two-sample t-tests between temperature treatments. (e) The *RVE2* cold-switch in plants under-going different extents of cooling was examined using RT-qPCR, *left* FS(.1) isoform and *right* AE isoform. Plants were entrained in 20°C in LD cycles then subjected to the denoted cooling extent at the onset of dusk with samples harvested at time-points spanning mid-dusk to dawn. Data represents one experiment with pooled tissue from 9-13 plants per time-point. The DDCt method was used to calibrate normalised Ct values (using the average of Cts for IPP2 and ISU1 housekeeping genes) to the time-point demonstrating maximal expression (6h at 20 to 4°C for FS(.1), *left*; and 9h at 20°C for AE, *right*).

**Fig. S2. Persistent *RVE2* rhythms in free-run conditions.** Transcript isoform-specific expression of *RVE2* in constant light by RT-qPCR. (a) Using RT-qPCR the FS(.1) expression profile for plants sampled over 24h in LD at either 20°C or the equivalent time-points for cooling transition (20 to 4°C) was determined. The DDCt method was used to calibrate normalised Ct values (using the average of Cts for IPP2 and ISU1 housekeeping genes) to the time-point demonstrating maximal expression (6h time-point at ‘20 to 4°C’ – calibrated as 100%). Relative to this, FS(.1) at 20°C was estimated to be 15% (8h time-point). The equivalent comparison using RNA-seq data (Suppl Fig. S1A), estimates FS(.1) at 20°C at 9h to be 8% of the levels for the 6h time-point at ‘20 to 4°C’. Col-0 plants were released into free run constant light conditions at either 20°C or 12 °C after previous entrainment in DL at 20°C and the (b) FS(.1) and (c) AE (_ID2 and _JC4) expression levels monitored. Data is mean ±SEM (*n*=3) of normalised expression levels. For (b) the level of FS(.1) at −4h at 20°C was set to 15%. For (b) the −4h levels in DL were set to 15%, for (c) the AE levels at −4h at 20°C was set to 100%. (d) Period length (h) estimates for FS and AE rhythms in LL were determined using BRASS software for 48h of data (24 - 72h) of panels (b) and (c).

**Fig. S3. Altered expression of candidate target RVE2 genes in *rve2-2* during onset of cool nights.** (a) GABI-Kat T-DNA insertion line 508D10 in Col-0 (https://www.gabi-kat.de) maps to intron 1 of *RVE2*, the region of cold-switching AS. (b) Eleven T3 segregating lines (numbered #2 to #12) were genotyped using GABI-Kat primer set 1 (Supplementary **Table S1**) predicted to amplify wild-type (WT), expected size 929bp; lines #4, #6, #9, #10, #11 and #12 are candidate (-/-) homozygotes. (c) Each line was screened for sensitivity to sulfadiazine (SUL) antibiotic according to GAB-Kat guidelines. Around 40-50 seeds were plated alongside Col-0 on SUL-agar plates. Each line scored for sensitivity to SUL, representative regions of SUL-agar plates for each line presented with estimates of SUL sensitivity (% of seeds not germinating). Col-0 fails to germinate (100% sensitivity) on each of the SUL-agar plates. Lines #4 and #11 (green rectangles) are the most promising as -/- homozygotes (0% sensitivity). Other lines are presumably heterozygotes (+/−) apart from line #7 (red rectangle; 100% sensitivity) that is probably (+/+) homozygous. (d) Lines #4 and #11 (-/-) were selfed, and gDNA isolated from 3 independent sibling plants, genotyped using GABI-Kat primer set 1 (as above) and GABI-Kat primer set 5 (South-junction primers, Supplementary **Table S1**), predicted size approximately 650bp. Amplicons were co-electrophoresed with parental genotyping products. RT-PCR band patterns are consistent with both #4 and #11 lines being homozygous (-/-) mutants. Bulk seeds from these 3 sibling lines were pooled for #4 and #11 seed stocks. (e-j) Expression profiles of the *RVE2* isoforms in the *rve2-2*/Col-0 temperature and time series RNA-seq experiment were determined for the (e) .1 (FS) and _ID2 (AE), (f) _ID1, (g) _ID3, (h) _ID4, (i) _s1, and (j) _JC4 RVE2 isoforms. Data is TPM mean and +/− S.E.M, *n*=3, and data points at 120h are repeated from 96h. Alternating black-white rectangles denote 12 h dark-light phases, respectively. cooling onset from 20°C to 4°C initiated at 24h. Data shows absence of *RVE2* FS(.1) and AE (_ID2) – panel (f) - across the entire temperature and time series experiment, but rhythmic expression of other isoforms.

## Notes

### Competing Interest Statement

The authors have declared no competing interest.

