## Supplementary material for "*REVEILLE2* Thermosensitive Splicing: A Molecular Basis for the Integration of Nocturnal Temperature Information by the Arabidopsis Circadian Clock": Suppl.

### Supplemental Figure 1

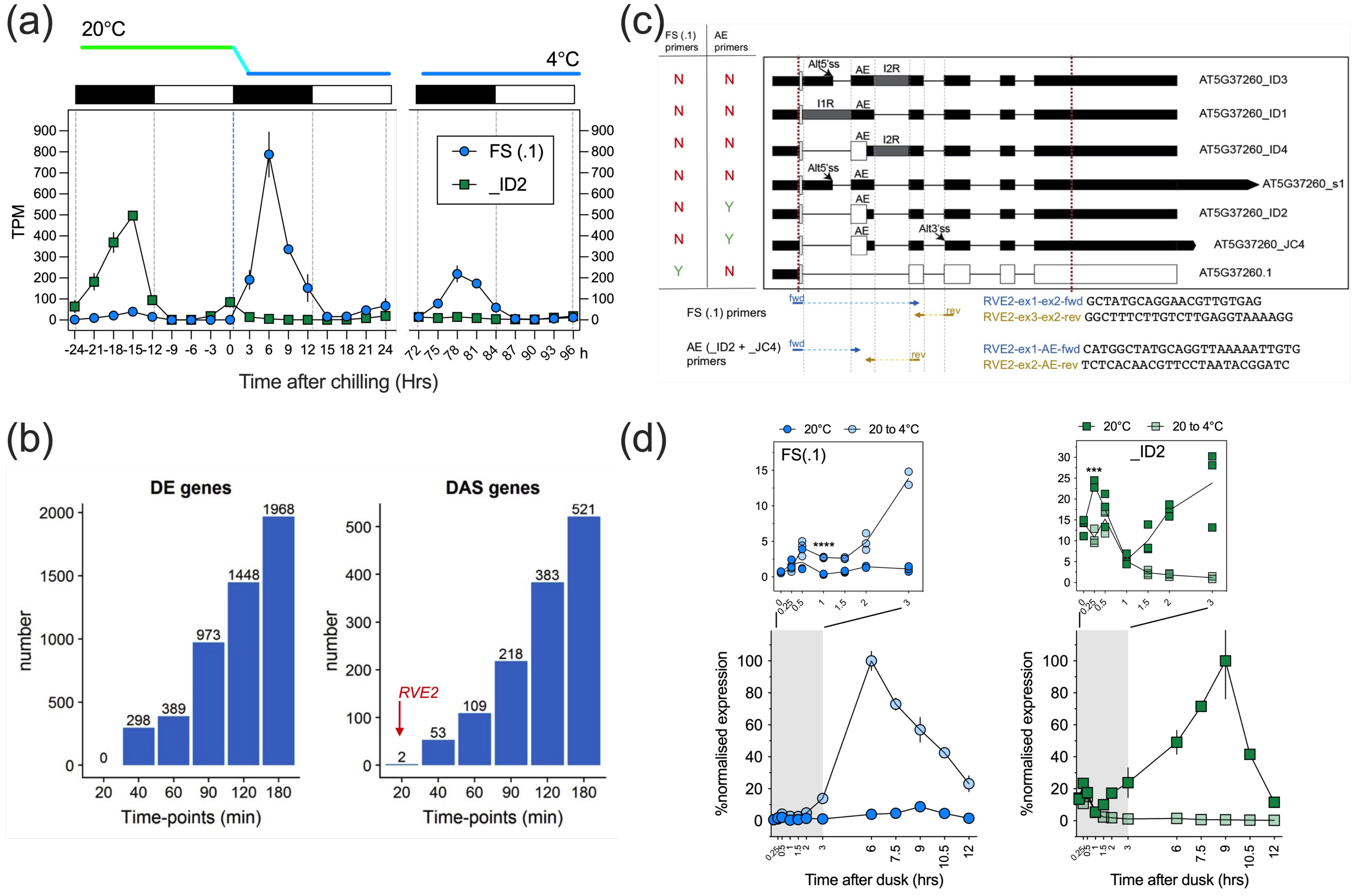

**Fig. S1. The splicing driven *RVE2* cold switch**. (a) Simplified form of main Fig **1c** clearly showing the isoform switch of the two main *RVE2* isoforms, FS(.) and _ID2. Data of Calixto et al., 2018; mean and ± SEM (*n*=3) of normalised expression levels (TPM; Transcripts Per Million) for the denoted transcript isoforms, cooling from 24h (dashed blue vertical line), diurnal 12h dark/12h light conditions as black/white rectangles, respectively. (b) The speed of changes in expression/AS were examined for a high-resolution RNA-seq time-series across the first 3h of cooling (Supplementary Dataset **S2**). Plotting the time-point at which genes were first significantly DE or DAS (each gene is represented only once) showed no significant DE genes at 20 min while two genes were significantly DAS at 20 min. One of two DAS genes with the most rapid significant AS was *RVE2*. (c) Graphical representation of predicted annealing locations of RT-qPCR primers (Supplementary **Table S1**) amplifying isoform specific *RVE2* amplicons. Primers spanned introns (dotted lines); forward (fwd), and reverse (rev) orientation denoted, primer sequence 5' to 3'. The FS(.1) primer combination is predicted to amplify FS(.1) specifically, whereas the AE primer combination is predicted to amplify both the _ID2 and _JC4 isoforms. (d) Expression profile of the two principal *RVE2* isoforms (FS(.1), *left* and AE (_ID2 and _JC4), *right*) using the RT-qPCR primers described in (c). Time points spanned 12h of dusk (at either 20ºC or cooling onset, 20 to 4ºC) except for closely spaced time points during the first 3h of dusk (yellow shaded area expanding to upper graphs). The $\Delta\Delta$Ct method was used to calibrate normalised Ct values (using the average of Cts for IPP2 and ISU1 housekeeping genes) to the time-point demonstrating maximal expression (6h at 20 to 4ºC for FS(.1), *left*; and 9h at 20ºC for AE, *right*). Data is mean ±SEM (*n*=3) of normalised expression levels. Statistical summaries (*** = $p \leq 0.001$, **** = $p \leq 0.0001$) for paired two-sample t-tests between temperature treatments.

### Supplemental Figure 2

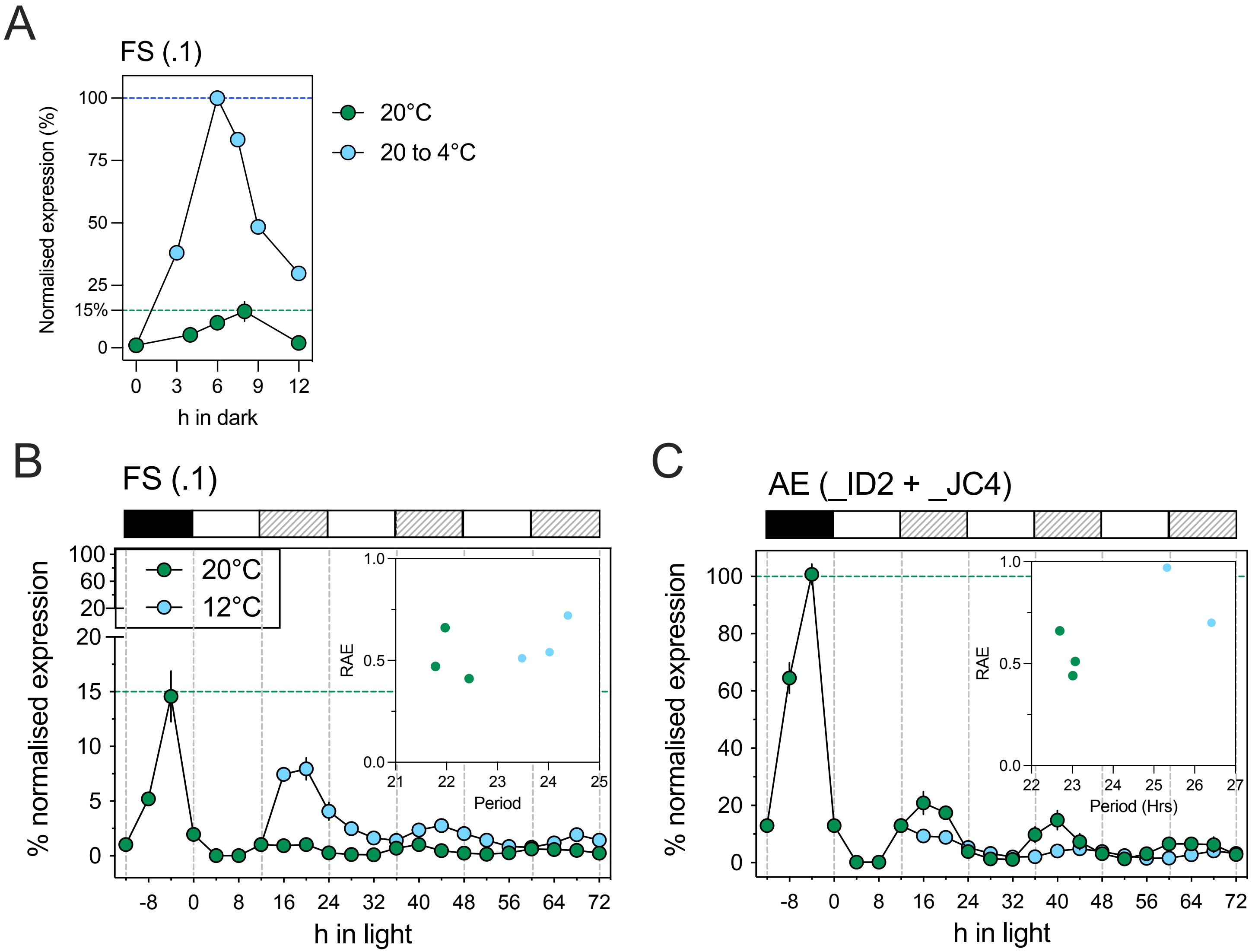


**Fig. S2. Persistent *RVE2* rhythms in free-run conditions.** Transcript isoform-specific expression of *RVE2* in constant light by RT-qPCR. (a) Using RT-qPCR the FS(.1) expression profile for plants sampled over 24h in LD at either 20ºC or the equivalent time-points for cooling transition (20 to 4ºC) was determined. The ΔΔCt method was used to calibrate normalised Ct values (using the average of Cts for *IPP2* and *ISU1* housekeeping genes) to the time-point demonstrating maximal expression (6h time-point at '20 to 4ºC' – calibrated as 100%). Relative to this, FS(.1) at 20ºC was estimated to be 15% (8h time-point). The equivalent comparison using RNA-seq data (Suppl Fig. S1A), estimates FS(.1) at 20ºC at 9h to be 8% of the levels for the 6h time-point at '20 to 4ºC'. Col-0 plants were released into free run constant light conditions at either 20ºC or 12 ºC after previous entrainment in DL at 20ºC and the (b) FS(.1) and (c) AE (_ID2 and _JC4) expression levels monitored. Data is mean ±SEM (*n*=3) of normalised expression levels. For (b) the level of FS(.1) at -4h at 20ºC was set to 15%, for (c) the AE levels at -4h at 20ºC was set to 100%. A comparison of period estimates against Relative Amplitude Error (RAE) is inset into (b) and (c) for each of the biological repeats.

### Supplemental Figure 3

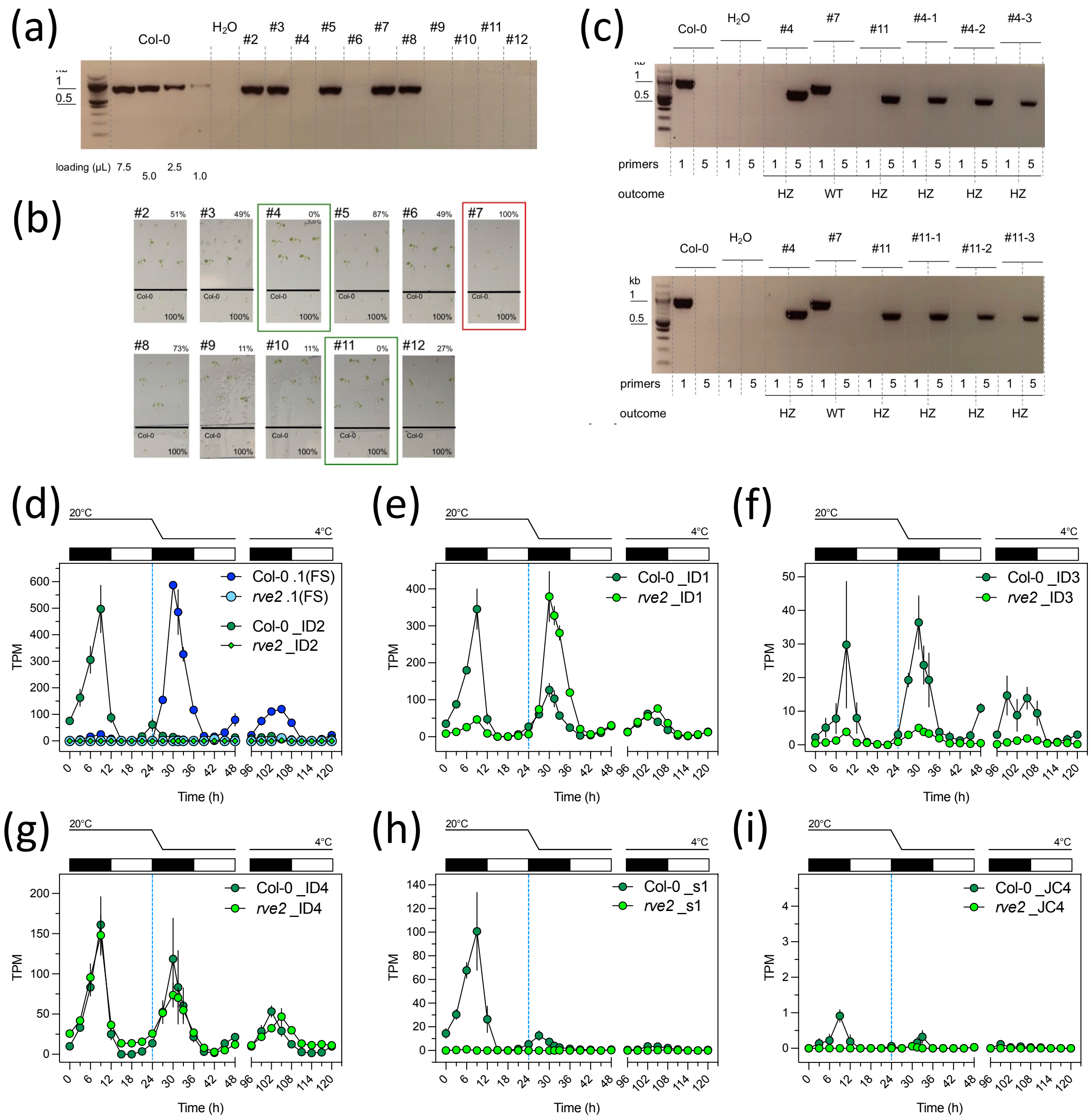


**Fig. S3. Altered expression of *RVE2* transcript isoforms in *rve2-2* during onset of cool nights. (a)** Eleven T3 segregating lines (numbered #2 to #12) were genotyped using GABI-Kat primer set 1 (Supplementary **Table S1**) predicted to amplify wild-type (WT), expected size 929bp; lines #4, #6, #9, #10, #11 and #12 are candidate (-/-) homozygotes. **(b)** Each line was screened for sensitivity to sulfadiazine (SUL) antibiotic according to GAB-Kat guidelines. Around 40-50 seeds were plated alongside Col-0 on SUL-agar plates. Each line scored for sensitivity to SUL, representative regions of SUL-agar plates for each line presented with estimates of SUL sensitivity (% of seeds not germinating). Col-0 fails to germinate (100% sensitivity) on each of the SUL-agar plates. Lines #4 and #11 (green rectangles) are the most promising as -/- homozygotes (0% sensitivity). Other lines are presumably heterozygotes (+/-) apart from line #7 (red rectangle; 100% sensitivity) that is probably (+/+) homozygous. **(c)** Lines #4 and #11 (-/-) were selfed, and gDNA isolated from 3 independent sibling plants, genotyped using GABI-Kat primer set 1 (as above) and GABI-Kat primer set 5 (South-junction primers, Supplementary **Table S1**), predicted size approximately 650bp. Amplicons were co-electrophoresed with parental genotyping products. RT-PCR band patterns are consistent with both #4 and #11 lines being homozygous (-/-) mutants. Bulk seeds from these 3 sibling lines were pooled for #4 and #11 seed stocks. **(d-i)** Expression profiles of the *RVE2* isoforms in the *rve2-2*/Col-0 temperature and time series RNA-seq experiment were determined for (E) .1 (FS) and _ID2 (AE), (F) _ID1, (G) _ID3, (H) _ID4, (I) _s1, and (J) _JC4 RVE2 isoforms. Data is TPM mean and +/- S.E.M, *n*=3, and data points at 120h are repeated from 96h. Alternating black-white rectangles denote 12 h dark-light phases, respectively. cooling onset from 20ºC to 4ºC initiated at 24h.
